# Arl15 upregulates the TGFβ family signaling by promoting the assembly of the Smad-complex

**DOI:** 10.1101/2021.12.14.472727

**Authors:** Meng Shi, Hieng Chiong Tie, Divyanshu Mahajan, Xiuping Sun, Yan Zhou, Boon Kim Boh, Leah A. Vardy, Lei Lu

## Abstract

The hallmark event of the canonical transforming growth factor β (TGFβ) family signaling is the assembly of the Smad-complex, consisting of the common Smad, Smad4, and phosphorylated receptor-regulated Smads. How the Smad-complex is assembled and regulated is still unclear. Here, we report that active Arl15, an Arf-like small G protein, specifically binds to the MH2 domain of Smad4 and colocalizes with Smad4 at the endolysosome. The binding relieves the autoinhibition of Smad4, which is imposed by the intramolecular interaction between its MH1 and MH2 domains. Activated Smad4 subsequently interacts with phosphorylated receptor-regulated Smads, forming the Smad-complex. Our observations suggest that Smad4 functions as an effector and a GTPase activating protein (GAP) of Arl15. Assembly of the Smad-complex enhances the GAP activity of Smad4 toward Arl15, therefore dissociating Arl15 before the nuclear translocation of the Smad-complex. Our data further demonstrate that Arl15 positively regulates the TGFβ family signaling.

## Introduction

The transforming growth factor β (TGFβ) family signaling pathway is initiated by the TGFβ family cytokines, consisting of TGFβs, nodals, activins, growth and differentiation factors, and bone morphogenetic proteins (BMPs)(Derynck and Budi, 2019; Massague, 2012; Schmierer and Hill, 2007; Wrana, 2013). The pathway can profoundly affect the development and homeostasis of animal tissue and has long been recognized as one of the most critical contributors to multiple human diseases such as cancers, fibrotic disorders, and cardiovascular diseases(Goumans and Ten Dijke, 2018; Kim et al., 2018; Seoane and Gomis, 2017). The core molecular framework of this pathway was outlined more than a decade ago(Massague, 2012; Schmierer and Hill, 2007). In the canonical TGFβ and BMP signaling pathways (hereafter referred to as the TGFβ and BMP signaling pathways), the TGFβ family cytokines first bind to and activate the type II and I receptor kinases, which phosphorylate receptor-regulated Smads (R-Smads), including TGFβ-specific Smad2 and 3 (hereafter referred to as TGFβ R-Smads) and BMP-specific Smad1, 5 and 8 (hereafter referred to as BMP R-Smads). Smads share a three-domain organization comprising N-terminal MH1 and C-terminal MH2 domain connected by a linker region. The type I receptor kinase phosphorylates the MH2 domain of an R-Smad. After phosphorylation, phospho-R-Smads and Smad4 assemble a complex, usually a heterotrimer with two phospho-R-Smads and one Smad4, via the association of their MH2 domains(Chacko et al., 2001; Chacko et al., 2004). Eventually, the Smad-complex translocates to the nucleus and executes genomic actions by chromatin remodeling, transcriptional activation, or repression of responsive genes. The formation of phospho-R-Smad and Smad4 complex is a central event in the TGFβ family signaling pathway. However, we still do not entirely understand this event at the molecular and cellular level. A fundamental question is how the assembly of the Smad-complex is initiated and regulated. Hata *et al*. previously reported that the self-autoinhibition of Smads regulates the assembly(Hata et al., 1997). They found that, in Smad4, the MH1 domain intramolecularly interacts with the MH2 domain, therefore preventing full-length Smad4 from interacting with R-Smads. What relieves the autoinhibition of Smad4 and how the resulting active Smad4 subsequently complexes with R-Smads are currently unknown.

Small G proteins are molecular switches that regulate diverse cellular processes. In the G protein cycle(Cherfils and Zeghouf, 2013), the guanine nucleotide exchange factor (GEF) activates an inactive or GDP-bound G protein to become the GTP-loaded or active form, which binds to its effectors and triggers cellular effects. The active G protein requires the GTPase activating protein (GAP) to hydrolyze the bound GTP and terminate its active state, completing the G protein cycle by returning it to its initial GDP-bound or inactive form. Arf-family G proteins are divided into Arf, Sar, and Arf-like (Arl) groups(Donaldson and Jackson, 2011; Sztul et al., 2019). Arf and Sar group members have well-documented cellular roles in recruiting vesicular coats and regulating lipid production. Arl group has the most members (>20), but most of them are poorly studied. They have diverse cellular functions, such as membrane trafficking, organelle positioning, microtubule dynamics, and ciliogenesis(Donaldson and Jackson, 2011; Sztul et al., 2019).

Arl15 is an uncharacterized Arl group member. It becomes interesting after a series of genome-wide association studies linked its gene locus to rheumatoid arthritis, multiple metabolic traits, such as body shape, blood lipid level, and magnesium homeostasis, and metabolic diseases, such as type 2 diabetes mellitus, coronary heart disease, and childhood obesity(Corre et al., 2018; Danila et al., 2013; Glessner et al., 2010; Li et al., 2014; Negi et al., 2013; Replication et al., 2014; Richards et al., 2009; Ried et al., 2016; Sun et al., 2015; Willer et al., 2013). It is unclear whether *ARL15* is the causative gene and how its genetic changes can lead to the aforementioned diseases since its molecular and cellular functions are largely unknown. Recent studies suggested that Arl15 might play a role in insulin signaling, adiponectin secretion, and adipogenesis(Rocha et al., 2017; Zhao et al., 2017). Here, we identified Arl15 as a novel regulator of the TGFβ family signaling. We found that it can directly bind to and activate autoinhibited Smad4 to promote the assembly of the Smad-complex. At the same time, the Smad4-containing complex negatively feedbacks to Arl15 by accelerating the GTP hydrolysis of Arl15. Therefore, our data demonstrate that Smad4 acts as an effector and GAP for small G protein Arl15.

## Results

### Arl15-GTP can directly interact with Smad4

During our systematic study of Arl group small G proteins, we focused on Arl15 due to its genetic implication in human diseases(Corre et al., 2018; Danila et al., 2013; Glessner et al., 2010; Li et al., 2014; Negi et al., 2013; Replication et al., 2014; Richards et al., 2009; Ried et al., 2016; Sun et al., 2015; Willer et al., 2013). Arl15 is ubiquitously expressed in human tissues, and its orthologs are present in most metazoans (Supplementary Fig. 1a). We found that exogenously expressed and endogenous Arl15 localized to the Golgi (Fig. 1a; Supplementary. Fig. 1b). Similar to other small G proteins(Feig, 1999; Sztul et al., 2019), GTP non-hydrolyzable mutation (GTP-bound or active form mutation), A86L (hereafter referred to as AL; see below for the experimental confirmation), and GDP-bound or inactive form mutation, T46N (hereafter referred to as TN), were introduced to Arl15. We found that both mutants localized to the Golgi similar to the wild type (WT) (Supplementary Fig. 1b). In addition to the Golgi, WT and AL-mutant Arl15-GFP were also detected at the plasma membrane (PM), early endosome (EE), late endosome (LE), and lysosome (Fig. 1b; Supplementary Fig. 1c).

**Figure 1.**
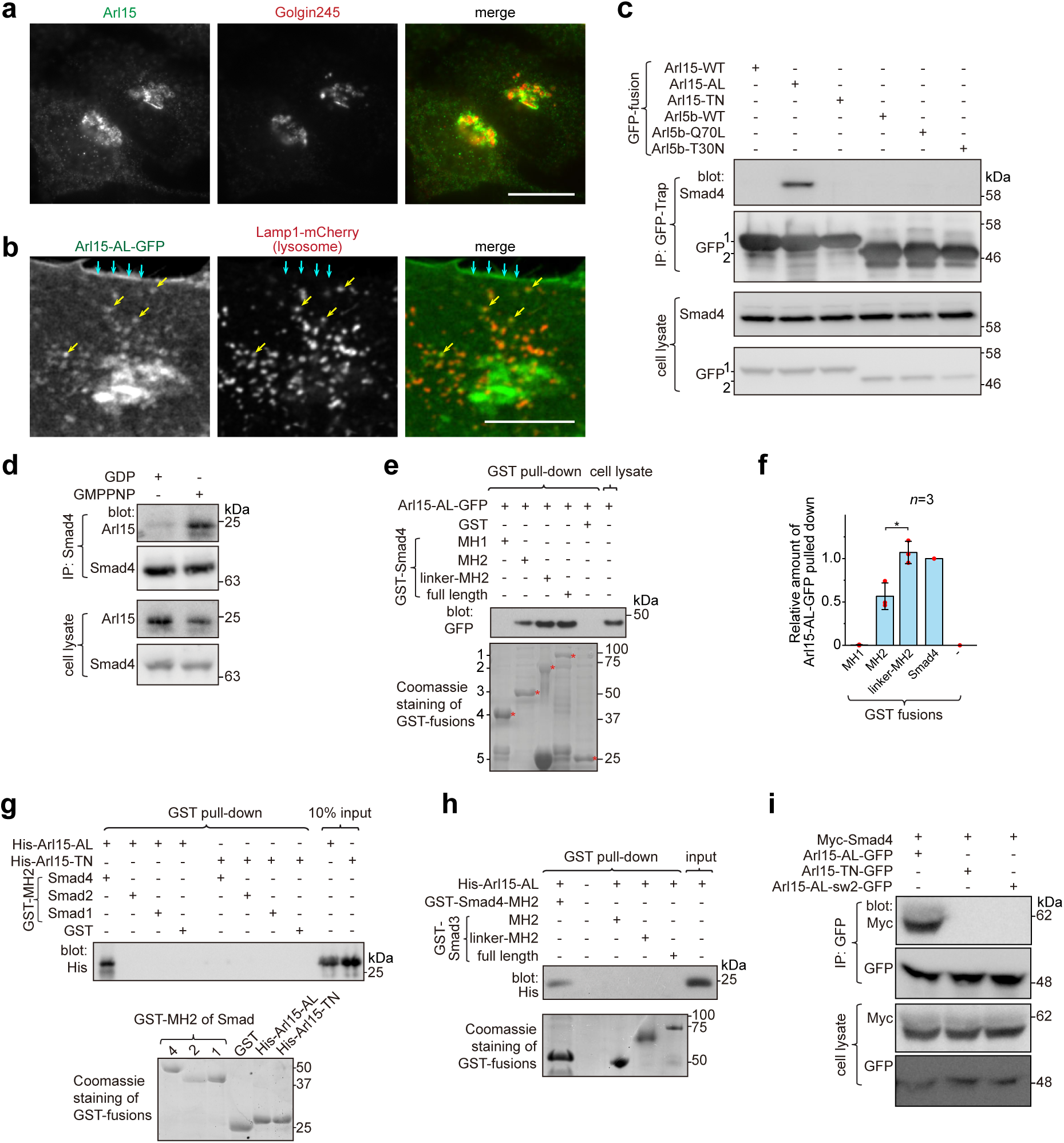
Arl15-GTP interacts with Smad4. **a** Endogenous Arl15 and Golgin245 were stained by immunofluorescence in HeLa cells. Images were acquired by a wide-field microscope. **b** Arl15-AL-GFP localizes to the lysosome. Live HeLa cells transiently co-expressing Arl15-AL-GFP and Lamp1-mCherry were imaged under a confocal microscope. Yellow arrows, colocalized puncta; cyan arrows, PM. Scale bar, 10 µm. **c** The GTP-mutant form of Arl15 specifically co-IPed endogenous Smad4. HEK293T cell lysates transiently expressing C-terminally GFP-tagged small G proteins were incubated with GFP-Trap beads, and IPs were immunoblotted against Smad4, and GFP. 1 and 2 indicate Arl15-(WT, AL or TN)-GFP and Arl5b-(WT, Q70L or T30N)-GFP bands, respectively. Arl5b serves as a negative control. **d** Endogenous Smad4 co-IPed Arl15 in the presence of GMPPNP, but not GDP. HEK293T cell lysates were incubated with anti-Smad4 antibody in the presence of 1 µM GMPPNP or GDP, and IPs were immunoblotted against Arl15 and Smad4. **e**,**f** The Smad4-MH2 domain specifically pulled down the GTP-mutant form of Arl15. In **e**, bead-immobilized GST-fusions of Smad4 fragments were incubated with HEK293T cell lysate expressing Arl15-AL-GFP, and pull-downs were immunoblotted against GFP. The loading of GST-fusions was shown below by Coomassie staining. 1, GST-Smad4; 2, GST-Smad4-linker-MH2; 3, GST-Smad4-MH2, 4, GST-Smad4-MH1 and 5, GST. * indicates specific band. The immunoblot is quantified in **f**, in which the relative amount of Arl15-AL-GFP pulled down is calculated as the ratio of the intensity of the pull-down band to that of the cell lysate input band. Error bar, mean ± SD of *n* = 3 experiments. *p* values are from the *t*-test (unpaired and two-tailed). *, *p* ≤ 0.05. Red dot, individual data point. **g**, **h** MH2 domain of Smad4, but not that of Smad1, 2, and 3, directly interacts with the GTP-mutant form of Arl15. Bead-immobilized GST-fusions of MH2 domains were incubated with purified His-tagged Arl15-AL or TN, and pull-downs were immunoblotted against His-Tag. Loading of fusion proteins was shown by Coomassie staining. **i** Switch-II region of Arl15 is required for its interaction with Smad4. HEK293T cell lysates expressing indicated proteins were incubated with GFP antibody, and IPs were immunoblotted against Myc-tag and GFP. In Arl15-AL-sw2-GFP, the switch-II region of Arl15-AL-GFP is replaced by that of Arl5b. Molecular weights (in kDa) are labeled in all immunoblots.

Yeast two-hybrid screening was subsequently performed to identify potential interacting partners of Arl15. Using Arl15-AL as a bait, we uncovered Smad4 as the most robust hit. Their interaction was confirmed by immunoprecipitation (IP) assays. We found that exogenously expressed and C-terminally GFP-tagged Arl15-AL, but not Arl15-TN, pulled down endogenous Smad4 (Fig. 1c). In contrast, Arl5b, another member of Arl group, showed negative results in either GTP (Q70L) or GDP-mutant (T30N) form(Shi et al., 2018). In the reverse IP, endogenous Smad4 pulled down substantially more endogenous Arl15 in the presence of guanosine 5′-[β,γ-imido]triphosphate (GMPPNP), a non-hydrolyzable GTP analog, than GDP (Fig. 1d). In Figure 1d, the weak pull-down band in the GDP panel is likely due to the cellular GTP. To explore which domain or region of Smad4 interacts with active Arl15, we prepared GST-fused Smad4 fragments (Supplementary Fig. 2a) and tested if they pull down Arl15-AL-GFP expressed in cell lysate (Fig. 1e; Supplementary. Fig. 2b). We found that the Smad4-MH2 domain, but not the MH1 domain or linker region, is sufficient to interact with Arl15-GTP. We noticed that the addition of the linker region significantly increased the pull-down of Arl15-GTP by the MH2 domain, as shown in pull-downs by GST-tagged Smad4-linker-MH2 and full-length Smad4 (Fig. 1e,f), demonstrating that the linker region probably contributes to the interaction too.

Although all MH2 domains share a similarity in sequences and structures, using purified GST-fusion proteins, we found that, while none of these MH2 domains interacted with His-Arl15-TN (Fig. 1g), only the MH2 domain of Smad4, but not that of Smad1, 2, and 3, directly bound to purified His-Arl15-AL (Fig. 1g,h). Extending the MH2 domain of Smad3 to include its linker region did not make the resulting chimera, GST-Smad3-linker-MH2, interact with Arl15-AL either (Fig. 1h). Since MH2 domains of BMP R-Smads, including Smad1, 5, and 8, share almost identical sequences with > 90% identity, our findings indicate that Arl15-GTP directly interacts with Smad4, but not TGFβ and BMP R-Smads.

Many interactions between a small G protein and its effectors involve the switch-II region of the small G protein, which undergoes disorder-to-order transition upon GTP-binding(Cherfils and Zeghouf, 2013; Vetter and Wittinghofer, 2001). To investigate the role of the switch-II region, we further mutated Arl15-AL by swapping its switch-II region with that of Arl5b, another Arl group small G protein that localizes to the Golgi(Shi et al., 2018) (Supplementary Fig. 1a). The resulting mutant, Arl15-AL-sw2, was able to bind to GTP as demonstrated by the GTP-agarose pull-down assay (Supplementary Fig. 2c), suggesting that the mutant might fold properly. In the subsequent Co-IP, we found that Arl15-AL-sw2-GFP failed to interact with co-expressed Myc-Smad4 (Fig. 1i), confirming the essential role of the switch-II in the interaction between Arl15 and Smad4. In summary, our data demonstrate that Arl15-GTP can directly and specifically interact with the MH2 domain of Smad4.

### Arl15-GTP colocalizes with Smad4 at the endolysosome

Smads likely localize to the endosome since they interact with SARA and endofin, adaptor proteins that possess endosome-targeting FYVE domains (Chen et al., 2007; Gillooly et al., 2001; Seet and Hong, 2001; Shi et al., 2007; Tsukazaki et al., 1998). Although immunostaining did not reveal a clear membrane association of Smad4, our live-cell confocal imaging uncovered a limited colocalization between mCherry-Smad4 and Arl15-AL-GFP at punctate structures (Fig. 2a). The weak punctate appearance of mCherry-Smad4 is likely due to the masking effect of its high cytosolic concentration. Alternatively, it might be due to the closed conformation of Smad4, which is formed by the intramolecular interaction between its MH1 and MH2 domains (see below). Therefore, we tested the mCherry-tagged Smad4-MH2 domain, which does not possess the inhibitory MH1 domain (Supplementary Fig. 2a). We observed that it displayed a much more robust punctate pattern, which colocalized with Arl15-AL-GFP (Fig. 2b,c). Further study revealed that these Smad4-MH2 positive puncta are primarily the EE, LE, and lysosome but not the recycling endosome (RE) (Fig. 2c,d). Together with our study of Arl15 (Fig. 1b; Supplementary Fig. 1c), our imaging data suggest that Arl15-GTP might interact with Smad4 at the endolysosome. The absence of the Smad4-MH2 domain at the Golgi, where most Arl15-AL-GFP resides, suggests an unknown *in vivo* mechanism restricting their interaction to the endolysosomal membrane.

**Figure 2.**
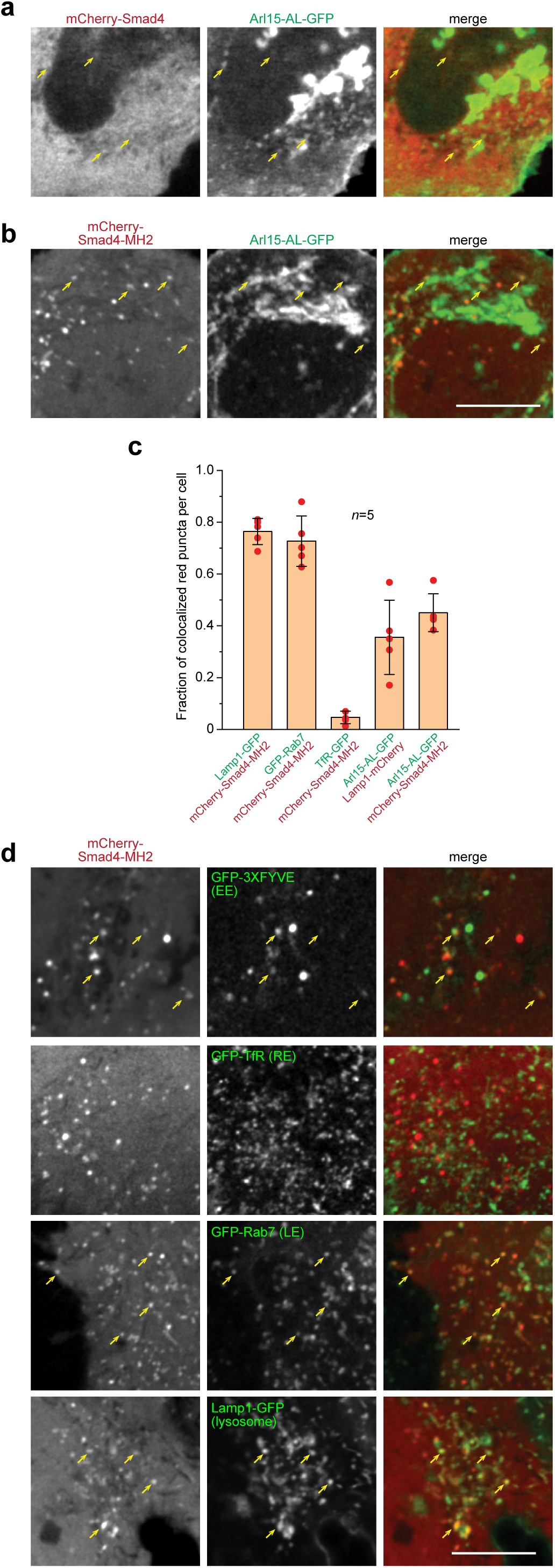
Smad4 colocalizes with the GTP-mutant form of Arl15 at the endolysosome. HeLa cells under the normal culture condition were used. **a**,**b** Smad4-MH2 displays better colocalization with the GTP-mutant form of Arl15 than full-length Smad4 at the endolysosome. Live HeLa cells co-expressing indicated mCherry and GFP-tagged proteins were imaged under a confocal microscope. Note that they colocalize at the endolysosome but not the Golgi. **c** Quantitative colocalization between Smad4-MH2 and various endolysosome markers. *n* = 5 cells were imaged, and all red puncta (mCherry-Smad4-MH2 or Lamp1-mCherry) within each image were examined. Fractions of red puncta that visually colocalize with green puncta were calculated and plotted. Error bar, mean ± SD (*n* = 5 cells). Red dot, individual data point. **d** Smad4-MH2 localizes to the EE, LE, and lysosome, but not the RE. Live HeLa cells co-expressing indicated mCherry and GFP-tagged proteins were imaged under the confocal microscope. GFP-tagged 3 × FYVE, TfR, Rab7, and Lamp1 are markers for the EE, RE, LE, and lysosome, respectively. Scale bar, 10 µm. Arrows indicate colocalization.

### Arl15-GTP indirectly interacts with R-Smads via Smad4

Although Arl15-GTP does not directly bind to R-Smads, we noticed that, in addition to Smad4, a significant amount of endogenous R-Smads, such as Smad2 and Smad1/5/8, was pulled down from cell lysates by GST-Arl15-AL, but not TN (Fig. 3a). The anti-Smad2/3 antibody used in Figure 3a should primarily detect endogenous Smad2 in our HEK293T cells since the band it detected was substantially reduced by the siRNA-mediated knockdown of Smad2, but not that of Smad3 (Supplementary Fig. 3a).

**Figure 3.**
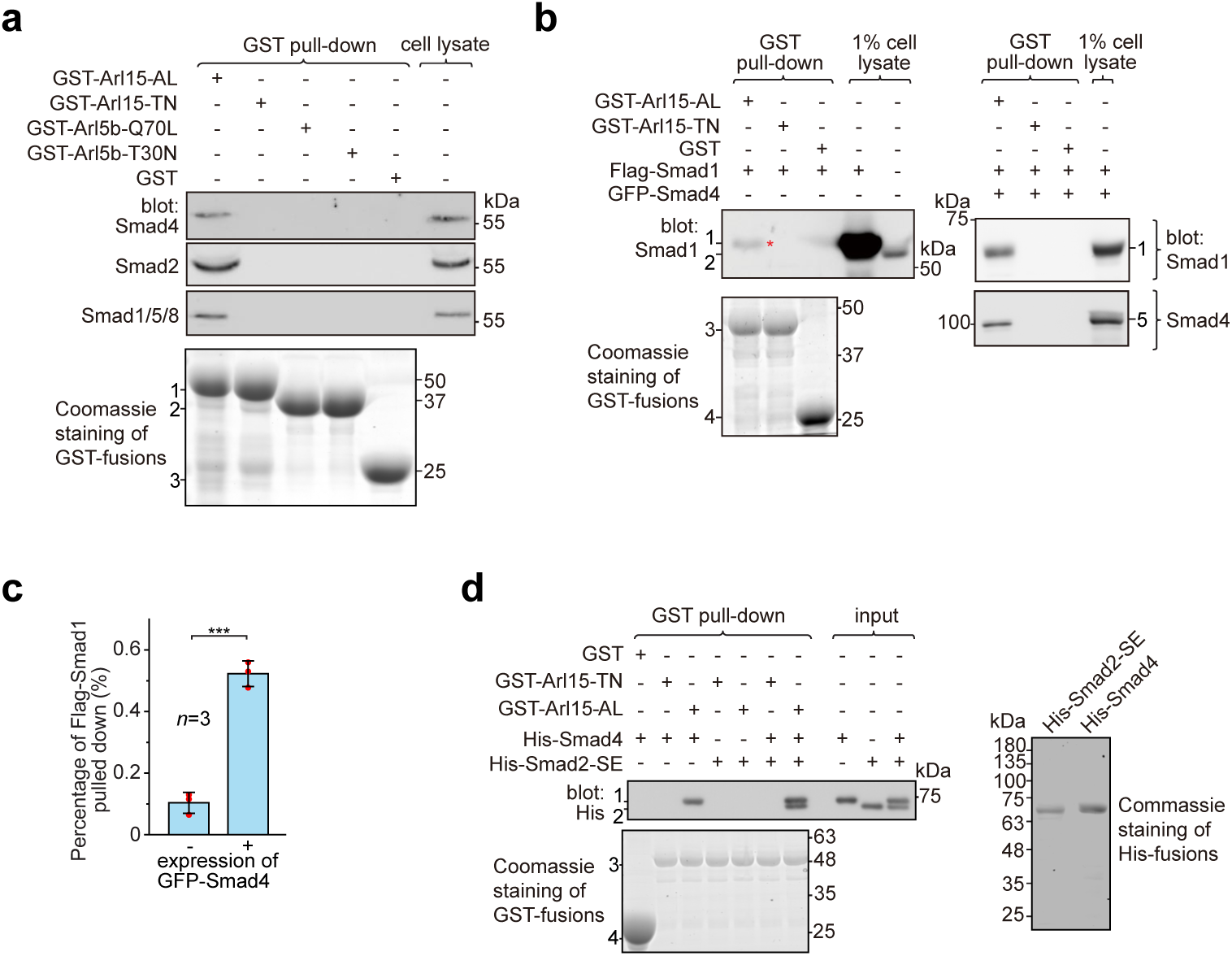
Arl15-GTP indirectly interacts with R-Smads via Smad4. HEK293T cells under the normal culture condition were used. **a** The GTP-mutant form of Arl15 specifically pulled down endogenous R-Smads in addition to Smad4. Bead-immobilized GST-fusion proteins were incubated with the cell lysate, and pull-downs and the cell lysate were immunoblotted against indicated Smads. 1, 2, and 3 indicate GST- Arl15 (AL or TN), GST-Arl5b (Q70L or T30N), and GST band. Arl5b serves as a negative control. **b**, **c** The GTP-mutant form of Arl15 pulled down more exogenously expressed Smad1 when Smad4 was co-expressed. In **b**, bead-immobilized GST-fusion proteins were incubated with cell lysates expressing indicated proteins, and pull-downs and the cell lysates were immunoblotted against Smad1 and 4. 1, Flag-Smad1; 2, endogenous Smad1/5/8; 3, GST-Arl15 (AL or TN); 4, GST; 5, GFP-Smad4; *, the weak band of Flag-Smad1 that was pulled down without co-expression of Smad4. Percentage of Flag-Smad1 pulled down, calculated as the ratio of the intensity of the pull-down band to that of the corresponding 1% cell lysate input band, is plotted in **c**. Error bar, mean ± SD of *n* = 3 experiments. *p* values are from the *t*-test (unpaired and two-tailed). ***, *p* ≤ 0.0005. Red dot, individual data point. **d** Arl15-GTP, Smad4 and Smad2 can assemble into a complex. Bead-immobilized GST-fusion proteins were incubated with indicated purified His-tagged Smads, and pull-downs were immunoblotted against His-tag. 1, His-Smad4; 2, His-Smad2-SE; 3, GST-Arl15 (AL or TN); 4, GST. The loading of fusion proteins is shown by Coomassie staining in **a**, **b** and **d**. Molecular weights (in kDa) are labeled in all immunoblots and gels.

Under normal cell culture condition, TGFβs in the serum(Danielpour et al., 1989) initiate a basal level of the TGFβ signaling to phosphorylate R-Smads. Since Smad4 can form a complex with phospho-R-Smads via their MH2 domains(Chacko et al., 2001; Chacko et al., 2004), our finding suggests that Smad4 probably bridges the indirect interaction between Arl15-GTP and phospho-R-Smads. The hypothesis was subsequently confirmed by the observation that bead-immobilized GST-Arl15-AL retained substantially more exogenously expressed R-Smads such as Smad1 and 2 (the BMP and TGFβ R-Smad respectively) in the presence than the absence of co-expressed Smad4 (Fig. 3b,c; Supplementary Fig. 3b). In GST-Arl15-AL pull-down without co-expressed Smad4, the residual amount of Smad1 and 2 (indicated by *) was likely due to the indirect interaction mediated by endogenous Smad4.

To avoid endogenous Smad4, we tested similar pull-downs using purified components (Fig. 3d). To mimic phosphorylated Smad2, we prepared His-Smad2 with S465E/S467E mutations (hereafter referred to as Smad2-SE)(Liu et al., 1997). We found that bead-immobilized GST-Arl15-AL pulled down Smad2-SE only in the presence of Smad4, therefore further supporting our hypothesis above. Likely, Arl15-GTP can indirectly interact with other R-Smads via Smad4 due to high identities shared among MH2 domains of TGFβ or BMP R-Smads. Hence, our data suggest that Arl15-GTP, R-Smad, and Smad4 might assemble as a complex.

### Arl15-GTP activates Smad4

Smads, including R-Smads and Smad4, adopt a closed conformation by an intramolecular association between MH1 and MH2 domains(Hata et al., 1997). Therefore, MH1 inhibits the corresponding MH2 domain and prevents the formation of the Smad-complex. We asked how Arl15-GTP affects the closed conformation of Smad4 by binding to its MH2 domain. Using truncated proteins comprising the MH1 or MH2 domain solely, we first confirmed that MH1 can interact with the corresponding MH2 domain in Smad4 and Smad2 (Fig. 4a-d). Furthermore, we observed that co-expressed Arl15-AL, but not Arl15-TN, substantially reduced the interaction between MH1 and MH2 domains of Smad4 (Fig. 4a,b), demonstrating that Arl15-GTP probably displaces Smad4-MH1 domain by interacting with Smad4-MH2 domain. Our results hence suggest that active Arl15 might open the closed conformation of Smad4.

**Figure 4.**
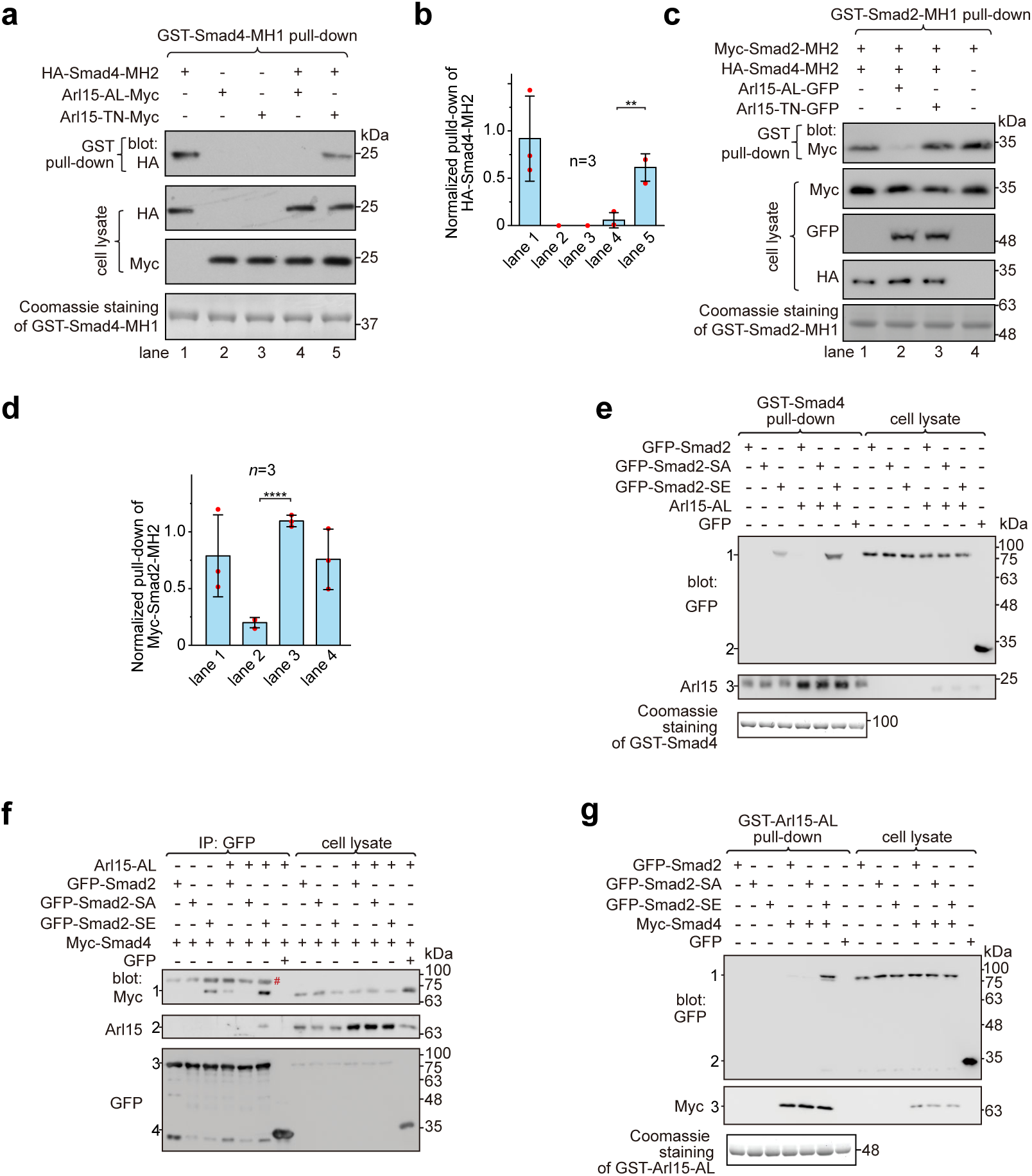
Arl15-GTP opens Smad4 and promotes assembly of the Smad-complex. HEK293T cells under the normal culture condition were used. **a** The GTP-mutant form of Arl15 opens Smad4 by inhibiting the intramolecular interaction between the MH1 and MH2 domain of Smad4. Bead-immobilized GST-Smad4-MH1 was incubated with the cell lysates expressing indicated proteins, and pull-downs and the cell lysates were immunoblotted against HA or Myc-tag. **b** The normalized pull-down of HA-Smad4-MH2 for assays conducted in **a**. The ratio of the intensity of the pull-down to that of the corresponding cell lysate band was calculated and plotted. **c** Arl15-GTP increases the intermolecular interaction between the Smad4-MH2 and Smad2-MH2 domain. Bead-immobilized GST-Smad2-MH1 domain was incubated with the cell lysates expressing indicated proteins, and pull-downs and the cell lysates were immunoblotted against indicated tags. **d** The normalized pull-down of Myc-Smad2-MH2 for assays conducted in **c**. Quantification was the same as in **b**. In **b**,**d**, error bar, mean ± SD of n = 3 experiments. *p* values are from the *t*-test (unpaired and two-tailed). **, *p* ≤ 0.005; ****, *p* ≤ 0.00005. Red dot, individual data point. **e**,**f** Arl15-GTP promotes the interaction between Smad4 and phospho-Smad2. In **e**, bead-immobilized GST-Smad4 was incubated with the cell lysates expressing indicated proteins, and pull-downs and the cell lysates were immunoblotted against indicated tags or protein. 1, GFP-Smad2 (WT, SA or SE); 2, GFP; 3, endogenous Arl15 or overexpressed Arl15-AL. In **f**, the cell lysates expressing indicated proteins were incubated with anti-GFP antibody, and co-IPs and the cell lysates were immunoblotted against indicated tags or protein. 1, Myc-Smad4; 2, endogenous Arl15 or overexpressed Arl15-AL; 3, GFP-Smad2 (WT, SA or SE); 4, GFP. #, non-specific band. **g** Arl15-GTP, Smad4, and phospho-Smad2 can assemble into a complex. Bead-immobilized GST-Arl15-AL was incubated with the cell lysates expressing indicated proteins, and pull-downs and the cell lysates were blotted. 1, GFP-Smad2 (WT, SA or SE); 2, GFP; 3, Myc-Smad4. In **a**,**c**,**e,** and **g**, loading of GST-fusion proteins is shown by Coomassie staining. Molecular weights (in kDa) are labeled in all immunoblots.

It has been documented that the Smad4-MH2 domain can interact with isolated MH2 domains of R-Smads in the absence of C-terminal phosphorylation(Hata et al., 1997; Wu et al., 2001). We found that the presence of the Smad4-MH2 domain did not substantially reduce the interaction between the MH1 and MH2 domains of Smad2 (Fig. 4c,d; compare lanes 1 and 4 of the first row of the gel blot). However, further addition of Arl15-AL (lane 2), but not Arl15-TN (lane 3), significantly weakened the interaction, suggesting that Arl15-GTP might promote the Smad4-MH2 domain to engage MH2 domains of R-Smads. Altogether, our biochemical data provide evidence that Arl15-GTP might activate Smad4 by relieving its MH2 domain from the intramolecular inhibition imposed by its MH1 domain.

### Arl15-GTP promotes the assembly of the Smad-complex

We next investigated the effect of Arl15-GTP on the assembly of the Smad-complex in the context of full-length Smads. In contrast to isolated MH2 domains, full-length R-Smads interact with Smad4 and assemble into a complex only after their C-termini are phosphorylated(Hata et al., 1997; Kretzschmar et al., 1997). Our data confirmed these reports and further revealed a molecular role of Arl15 in the assembly of the Smad-complex. First, bead-immobilized GST-Smad4 pulled down a substantial amount of Smad2-SE, but not WT and the non-phosphorylatable mutant, Smad2-S465A/S467A (hereafter referred to as Smad2-SA) (Fig. 4e); consistently, GFP-tagged Smad2-SE, but not WT or Smad2-SA, was found to interact with Myc-Smad4 in the co-IP assay (Fig. 4f). Second, only Smad2-SE, but not Smad2-SA, was pulled down by bead-immobilized Arl15-AL in a Smad4-dependent manner (Fig. 4g). Third and most importantly, we observed that the interaction between Smad2-SE and Smad4, that is, the formation of the Smad-complex, was substantially enhanced in the presence of Arl15-AL (Fig. 4e,f). A similar promoting effect of Arl15-AL on the interaction between Smad1, a BMP R-Smad, and Smad4 was also observed (Supplementary Fig. 4). Collectively, our results showed that Arl15-GTP might promote the assembly of the Smad-complex by binding to and activating the Smad4-MH2 domain.

### The Smad-complex functions as a GAP to inactivate Arl15-GTP

Once assembled in the TGFβ family signaling pathway, the Smad-complex enters the nucleus to initiate genomic actions(Derynck and Budi, 2019; Massague, 2012; Schmierer and Hill, 2007; Wrana, 2013). We then asked if Arl15-GTP co-translocates to the nucleus together with the Smad-complex. Our fluorescence imaging and nuclear fractionation did not detect Arl15 in the nucleus under TGFβ1 (Fig. 5a-c) or BMP2 treatment (Supplementary Fig. 5a). In contrast, phospho-Smad2/3 (Fig. 5a,b), phospho-Smad1/5/8 (Supplementary Fig. 5a), and Smad4 (Fig. 5b; Supplementary Fig. 5a) had increased presence in the nucleus under the same treatment, consistent with our current knowledge of the TGFβ family signaling pathway(Derynck and Budi, 2019; Massague, 2012; Schmierer and Hill, 2007; Wrana, 2013). Therefore, we reasoned that Arl15 might dissociate from the Smad-complex before the nuclear translocation of the latter. Our reasoning prompted us to test the hypothesis that the Smad-complex might act as a GAP to inactivate and consequently dissociate Arl15.

**Figure 5.**
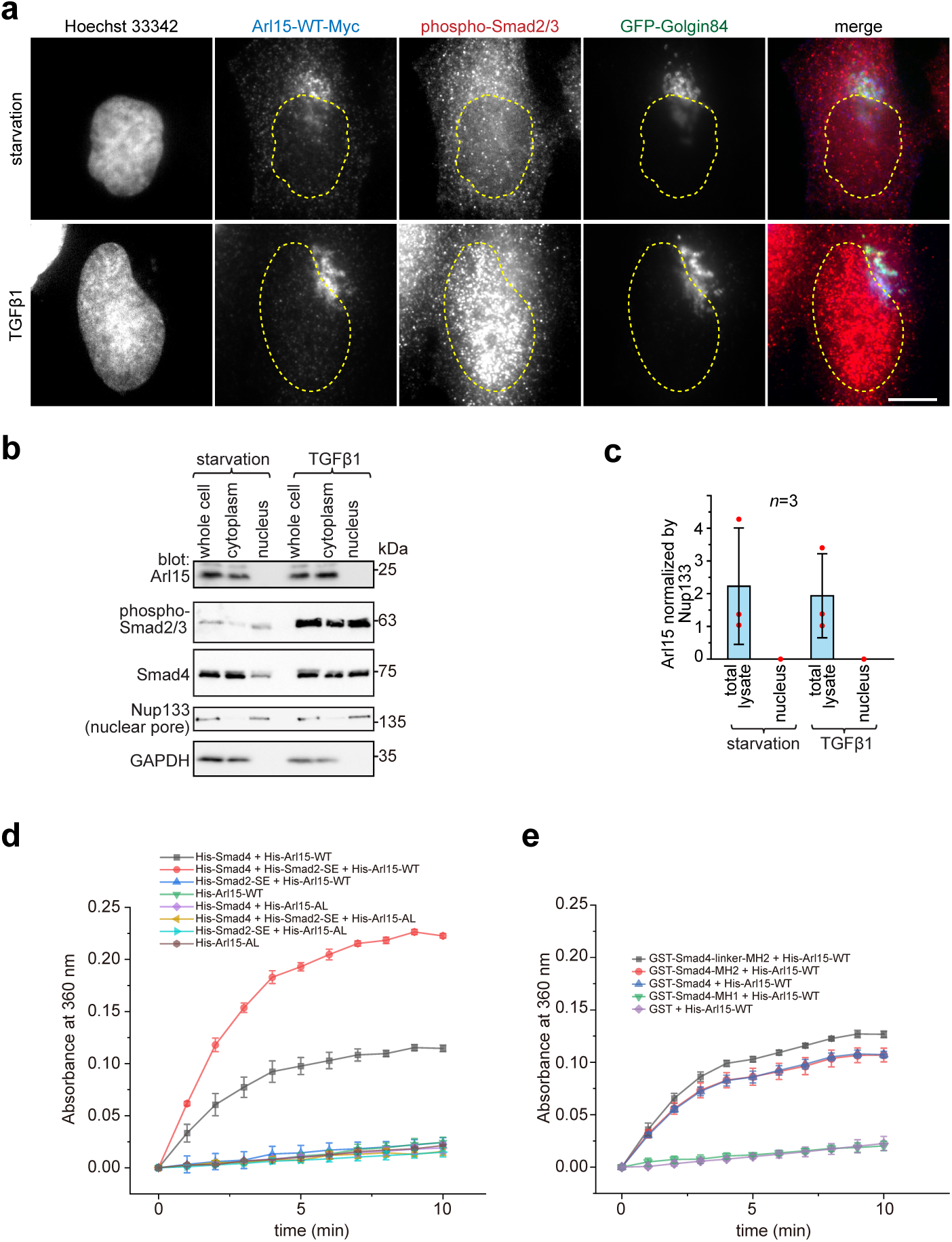
The Smad-complex functions as a GAP to inactivate and dissociate Arl15 before translocating to the nucleus. **a**-**c** Under TGFβ1 treatment, phospho-Smad2/3, but not Arl15, translocated to the nucleus. In **a**, serum-starved HeLa cells expressing GFP-Golgin84 (a Golgi marker) and Arl15-WT-Myc were either further serum-starved or treated with 10 ng ml^-1^ TGFβ1 for 1 h. Cells were stained for DNA (Hoechst 33342) and endogenous phospho-Smad2/3. Dotted line indicates contour of the nucleus. Scale bar, 10 µm. In **b**, serum-starved HeLa cells were either further serum-starved or treated with 10 ng ml^-1^ TGFβ1 for 1 h. Total cell lysate and nuclear and cytosol fractions were subjected to immunoblotting against indicated proteins. GAPDH, glyceraldehyde 3-phosphate dehydrogenase. Molecular weights (in kDa) are labeled in immunoblots. In **c**, assays conducted in **b** were quantified to show the relative amount of Arl15 normalized by Nup133. The ratio of the intensity of Arl15 band to that of the corresponding Nup133 band was calculated. Red dot, individual data point. **d** Phospho-Smad2 promotes the GAP activity of Smad4 toward Arl15. 40 µM GTP-loaded His-Arl15-WT or AL was incubated with 0.4 µM indicated His-Smads at 22 °C. Released inorganic phosphate was enzymatically converted and continuously monitored by absorbance at 360 nm. The absorbance was plotted against time. **e** The Smad4-MH2 domain possesses the GAP activity toward Arl15. The experiment was conducted as in **d**. GST-fused Smad4 fragments were used. In **c**, **d,** and **e**, error bar, mean ± SD of *n* = 3 experiments.

To that end, we first purified recombinant His-tagged Smad2-SE, Smad4, and Arl15 (WT or AL) (Supplementary Fig. 5b). Next, His-Arl15 (WT or AL) was first loaded with GTP and subsequently incubated with or without different combinations of His-Smad2-SE and His-Smad4. The inorganic phosphate released during the GTP hydrolysis was enzymatically converted and continuously monitored by spectrophotometry (see Materials and Methods). We found that Arl15-WT alone displayed a weak GTP-hydrolysis activity and that the GTP hydrolysis rate of Arl15 was substantially accelerated by the presence of Smad4 but not Smad2-SE, demonstrating Smad4 as a potential GAP for Arl15 (Fig. 5d). Interestingly, the addition of Smad2-SE greatly enhanced the GAP activity of Smad4 toward Arl15. As expected for a small G protein, we observed that AL mutation abolished the GTP hydrolysis activity of Arl15 in all cases — either alone or with the addition of Smad4 or Smad2-SE, retrospectively confirming AL as the GTPase-defective mutation. Using purified domains of Smad4 (Supplementary Fig. 5b), we mapped the GAP activity to the MH2 domain of Smad4 (Fig. 5e). The extension of the MH2 domain to include the linker region further increased the GAP activity (Fig. 5e). The positive effect of the linker region on the GAP activity of the MH2 domain is probably due to the enhanced interaction between the Smad4-MH2 domain and Arl15-GTP (Fig. 1e,f). In summary, our data suggest that, after the assembly of the Smad-complex, Smad4 might have an enhanced GAP activity and consequently inactivate Arl15 and dissociate the latter from the complex.

### Arl15-GTP is an essential and positive regulator of TGFβ and BMP signaling pathways

To understand the cellular significance of Arl15-Smad4 interaction, we investigated the effect of overexpression or depletion of Arl15 on TGFβ-induced transcriptions in HeLa cells. We first tested the transcription of Smad binding element×4-luc (SBE×4-luc), a luciferase reporter that comprises four tandem repeats of SBEs as its enhancer(Zawel et al., 1998). SBE×4-luc is commonly used for assaying the TGFβ R-Smad-dependent transcription or TGFβ signaling. We found that when HeLa cells were treated with a serum-free medium (serum-starvation or hereafter referred to as starvation), overexpression of Arl15-WT or AL, but not TN, was sufficient to stimulate the transcription of SBE×4-luc to ≥ 2-fold that of the control (Fig. 6a). However, the stimulation was much weaker than that of TGFβ1 treatment, which is ∼ 25-fold that of control (Fig. 6a). The stimulation was abolished by SB431542 (Supplementary Fig. 6a), a small molecule inhibitor of TGFβ type I receptor kinase activity(Laping et al., 2002). A possible explanation for these observations is that Arl15-AL might amplify the autocrine TGFβ signaling in HeLa cells(Qing et al., 2004) by promoting the formation of the Smad-complex. Similarly, we found that the overexpression of Arl15-WT or AL, but not TN, also stimulated the transcription of BRE-luc (Supplementary Fig. 6b), a luciferase reporter for assaying the BMP R-Smad-dependent transcription or BMP signaling(Korchynskyi and ten Dijke, 2002). On the other hand, when endogenous Arl15 was depleted by RNAi (Supplementary Fig. 6c), TGFβ1 or BMP2-induced reporter transcription was attenuated to less than half of the control, and the autocrine stimulated transcription also decreased under starvation (Fig. 6b; Supplementary Fig. 6d). Hence, our data imply that Arl15 is an essential and positive regulator of TGFβ and BMP signaling pathways.

**Figure 6.**
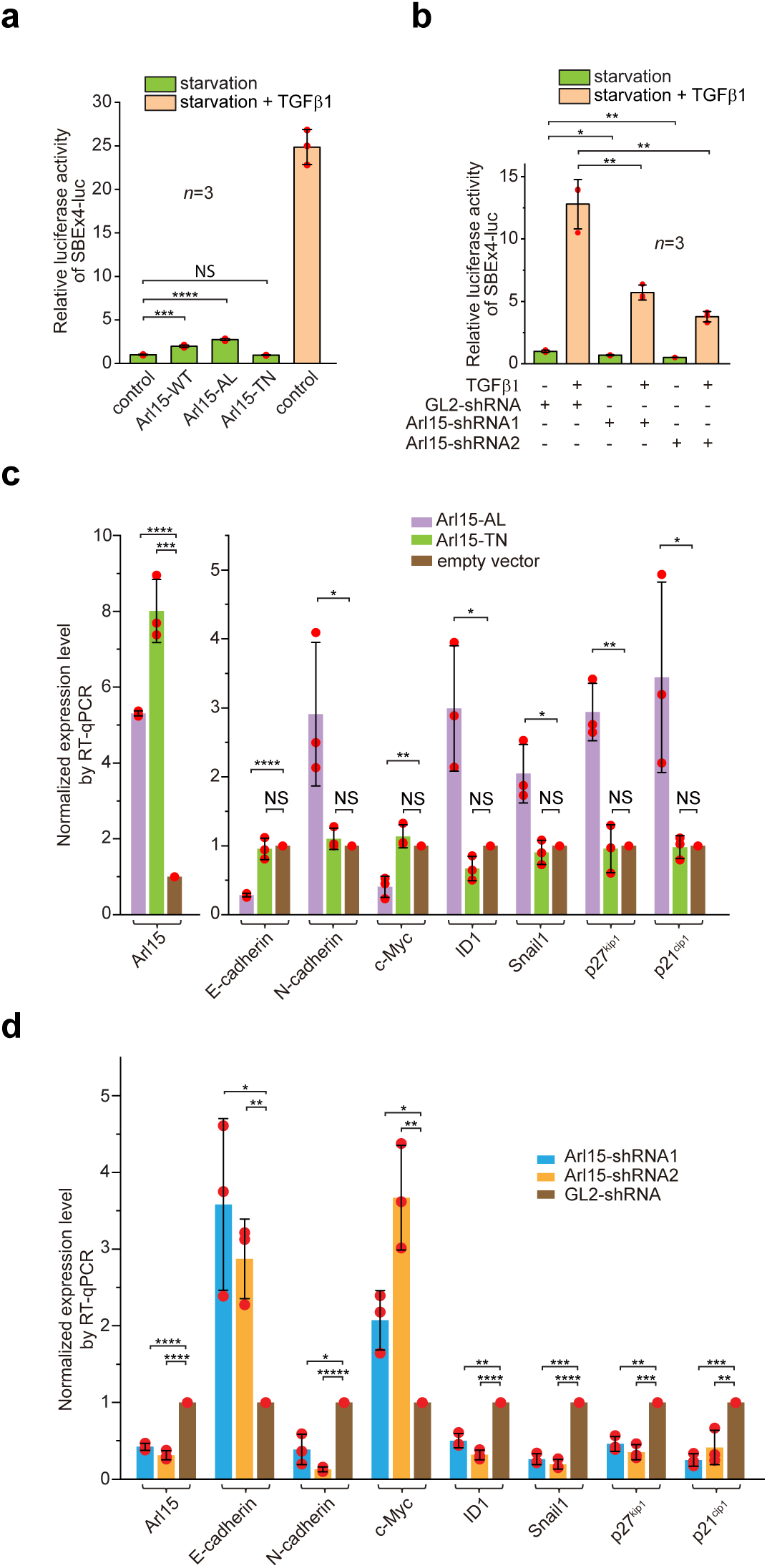
Arl15-GTP is an essential and positive regulator for the TGFβ family signaling pathway. **a** Arl15 can positively regulate the TGFβ signaling pathway since overexpressed Arl15-WT and AL, but not TN, promotes the transcription of SBE×4-luc reporter under starvation. HeLa cells co-expressing the SBE×4-driven firefly luciferase and SV40-driven renilla luciferase together with indicated Arl15 mutant or pBluescript SK vector DNA (control) were serum-starved for 24 h. For TGFβ1 induction, control cells were serum-starved for 4 h followed by 5 ng ml^-1^ TGFβ1 treatment for 20 h. Dual-luciferase assays were performed, and relative luciferase activities were subsequently acquired and normalized. **b** Arl15 is essential for efficient TGFβ1 signaling since its depletion reduces TGFβ1-stimulated transcription of SBE×4-luc reporter. After lentivirus-mediated knockdown of Arl15, HeLa cells co-expressing the dual-luciferase described in **a** were serum-starved for 4 h followed by either further starvation or 5 ng ml^-1^ TGFβ1 (starvation + TGFβ1) treatment for 20 h. Relative luciferase activities were subsequently acquired and normalized. GL2 is a non-targeting control shRNA. **c** Overexpression of Arl15-AL, but not TN, promotes the transcription of N-cadherin, ID1, Snail1, p27^kip1^, and p21^cip1^, and suppresses the transcription of E-cadherin and c-Myc. MCF7 cells were subjected to lentivirus-transduced overexpression of Arl15-AL or TN followed by starvation for 16 h. Transcripts of indicated genes were quantified by RT-qPCR and normalized by control (empty vector). **d** Opposite to overexpression, depletion of Arl15 suppresses TGFβ1-induced transcription of N-cadherin, ID1, Snail1, p27^kip1^, and p21^cip1^, and promotes the transcription of E-cadherin and c-Myc. After lentivirus-transduced knockdown of Arl15, MCF7 cells were subjected to 5 ng ml^-1^ TGFβ1 treatment for 72 h. Transcripts of indicated genes were quantified and normalized as in **c**. Error bar, mean ± SD of *n* = 3 experiments. *p* values were from the *t*-test (unpaired and two-tailed). NS, not significant (*p* > 0.05); *, *p* ≤ 0.05; **, *p* ≤ 0.005; ***, *p* ≤ 0.0005; ****, *p* ≤ 0.00005; *****, *p* ≤ 0.000005.

Since some Arl group small G proteins can regulate intracellular trafficking(Donaldson and Jackson, 2011; Sztul et al., 2019), we wondered if Arl15 indirectly regulates TGFβ family signaling by its role in intracellular trafficking. Therefore, we first investigated if Arl15 is required for secretory trafficking. Using ManII (a Golgi transmembrane glycosidase) and TNFα (a PM-targeted transmembrane protein) RUSH reporters(Boncompain et al., 2012), we found that Arl15 knockdown did not substantially affect the ER-to-Golgi (Supplementary Fig. 6e,f) and the subsequent Golgi-to-PM trafficking (Supplementary Fig. 6g,h). Next, we observed that TGFβ1-stimulated phosphorylation of Smad2/3 normally occurred upon knockdown of Arl15 (Supplementary Fig. 6i). Therefore, Arl15 is probably not required for steps leading to phosphorylation of Smad2/3, such as trafficking and maintenance of TGFβ1 receptors and Smad2/3. In summary, our data argue against a hypothesis that Arl15 indirectly regulates the TGFβ family signaling by intracellular trafficking. Instead, they support a model in which Arl15 regulates the TGFβ family signaling by promoting the assembly of the Smad-complex.

### Arl15 is essential to stimulate the transcription of TGFβ target genes

In addition to luciferase reporters, we also studied the effect of disrupting Arl15 on the cellular transcription profile of TGFβ target genes. To that end, we employed the MCF7 cell line, which represents early-stage or pre-malignant breast cancer cells(Comsa et al., 2015; Holliday and Speirs, 2011). In serum-starved MCF7 cells, overexpression of Arl15-AL, but not TN and empty vector control, was sufficient to upregulate the transcription of N-cadherin, ID1, Snail1, p27^kip1^, and p21^cip1^, and downregulate the transcription of E-cadherin and c-Myc (Fig. 6c). On the other hand, under the treatment of TGFβ1, depletion of Arl15 in MCF7 cells reversed the transcriptional trend of above genes compared to control knockdown, i.e., those that were upregulated by Arl15-AL overexpression became downregulated, and vice versa (Fig. 6d). A similar result was obtained when Arl15 was depleted in the MDA-MB-231 cell line, representing highly metastatic breast cancer cells (Supplementary Fig. 6j,k). Therefore, our observation correlates the activity of Arl15 with a gene transcription profile characteristic of the TGFβ-induced cytostasis and EMT in cancer cells(Hao et al., 2019; Lamouille et al., 2014; Seoane and Gomis, 2017).

Since the AL mutation compromises the GTP hydrolysis activity of Arl15, Arl15-AL should sequester the Smad-complex and inhibit the downstream transcription event. Indeed, the overexpression of Arl15-AL reduced the transcription of SBE×4-luc in HeLa cells under TGFβ1 treatment (Supplementary Fig. 6l). However, our luciferase assays indicated that overexpression of Arl15-AL upregulates both TGFβ and BMP pathways under starvation, although to a limited extent (Fig. 6a; Supplementary Fig. 6b). Therefore, we hypothesize that Arl15-AL might have two opposing effects on the Smad-complex – promoting its assembly and inhibiting its nuclear translocation. Under starvation, the assembly of the Smad-complex might be the rate-limiting step when cellular phospho-Smad2/3 concentration is low under autocrine stimulation. Arl15-AL might spontaneously dissociate from the Smad-complex, freeing the Smad-complex for its subsequent nuclear translocation. Hence, the net effect is that Arl15-AL promotes transcription of SBE×4-luc (Fig. 6a; Supplementary Fig. 6b). In contrast, under TGFβ1 treatment, when the cellular phospho-Smad2/3 concentration is high, the nuclear translocation of the Smad-complex becomes the rate-limiting step, and the net effect of Arl15-AL overexpression might be inhibitory.

### *in vitro* migration and invasion of cancerous cells require Arl15

The TGFβ signaling pathway is a well-known promoting factor for the metastasis of cancer cells(Hao et al., 2019; Lamouille et al., 2014; Seoane and Gomis, 2017). To explore the role of Arl15 in cancer metastasis, we assessed two key metastatic traits *in vitro*, cellular migration and invasion, using a highly metastatic breast cancer cell line – MDA-MB-231. By wound healing and collagen gel invasion assays, we observed that the migration and invasion of MDA-MB-231 cells were significantly reduced upon depletion of Arl15 (Fig. 7a-d), therefore demonstrating the indispensable role of Arl15 in the migration and invasion of metastatic cancerous cells *in vitro*. Although the precise role of Arl15 in tumorigenesis requires further investigation, our current data are consistent with what has been documented about the TGFβ signaling pathway in the metastasis of cancers(Derynck and Budi, 2019; Hao et al., 2019; Lamouille et al., 2014; Massague, 2012; Schmierer and Hill, 2007; Seoane and Gomis, 2017), thus further supporting Arl15 as an essential and positive regulator of this pathway.

**Figure 7.**
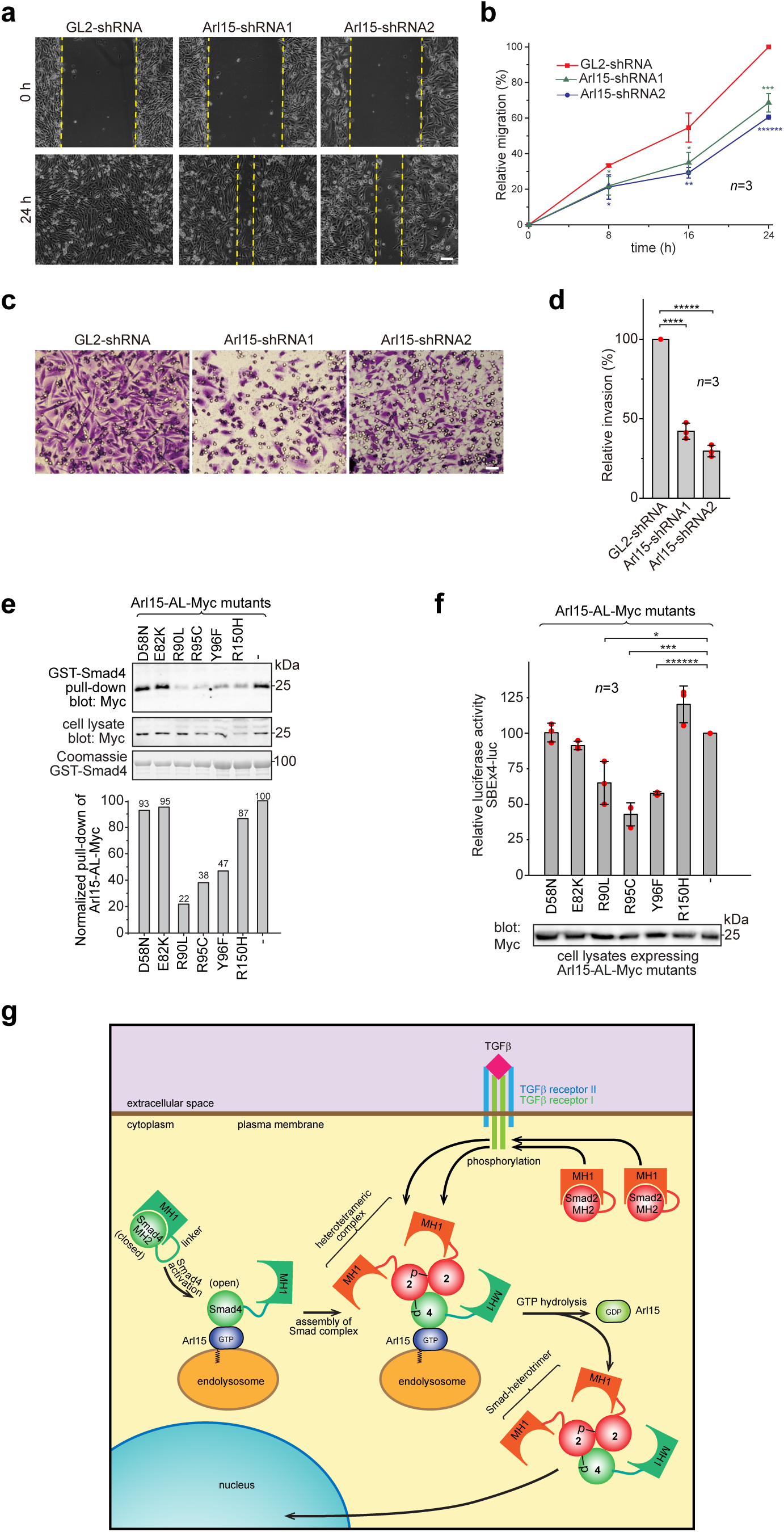
Implication of Arl15 in tumorigenesis by *in vitro* assays and mutation analysis, and our working model on how Arl15 regulates the TGFβ family signaling pathway. **a**, **b** Arl15 is required for *in vitro* migration of MDA-MB-231 cells. MDA-MB-231 cells were subjected to lentivirus-transduced knockdown using indicated shRNA. When cells reached confluency, a strip of cells was scraped off, and the resulting gap was live-imaged to monitor the migration of cells. The percentage of the relative migration (see Methods) is plotted in **b**. **c**,**d** Arl15 is required for *in vitro* invasion and migration of MDA-MB-231 cells. MDA-MB-231 cells were subjected to lentivirus-transduced knockdown using indicated shRNA and were subsequently placed into cell culture filter chambers with basement matrix. Cells that invaded through the matrix and migrated to the lower surface of the filter were stained in **c**. In **d**, the relative invasion was calculated as described in Materials and Methods. Scale bar, 100 µm. **e**,**f** Arl15 missense mutations identified from cancer patients compromise Arl15-Smad4 interaction and TGFβ signaling. In **e**, bead-immobilized GST-Smad4 was incubated with HeLa cell lysates expressing Arl15-AL-Myc harboring indicated mutation, and pull-downs and the cell lysates were immunoblotted against Myc-tag. Normalized pull-down of Arl15-AL-Myc was shown below, and it was calculated as the ratio of the intensity of the pull-down band to that of the corresponding cell lysate band. Loading of fusion proteins is shown by Coomassie staining. In **f**, HeLa cells were co-transfected to express the dual-luciferase and Arl15-AL-Myc with indicated mutation. After 20 h starvation, cells were subjected to the dual-luciferase assay and Western blot analysis for Myc-tag. In **b**, **d,** and **f**, error bar, mean ± SD of *n* = 3 experiments. *p* values were from the *t*-test (unpaired and two-tailed). *, *p* ≤ 0.05; ***, *p* ≤ 0.0005; ******, *p* ≤ 0.0000005. Red dot, individual data point. Molecular weights (in kDa) are labeled in all immunoblots. **g** A working model illustrating the molecular mechanism on how Arl15 regulates the TGFβ family signaling pathway. Smad2 is used as an example of the R-Smad. 2, Smad2; 4, Smad4; p, phosphate group at Smad2. See text for details.

### Some somatic mutations from cancer patients can compromise Arl15-Smad4 interaction

Using cancer genomic databases, we explored mutations in cancer patients that can disrupt Arl15-Smad4 interaction. We identified and tested somatic missense mutations at the G3-motif or switch-II region of Arl15: E82K, R90L, Y95C, and Y96F (Supplementary Fig. 1a) with two mutations at other locations, D58N and R150H. When each mutation was introduced in Arl15-AL, the resulting mutant protein displayed normal GTP-binding activity in the GTP-agarose pull-down assay (Supplementary Fig. 7), suggesting that these mutations might not disrupt the folding of Arl15. We found that the following single point mutations in the switch-II region specifically attenuated the Arl15-Smad4 interaction: R90L, Y95C, and Y96F, as demonstrated in our pull-down assays (Fig. 7e). These mutations also substantially reduced the stimulation of Arl15-AL in the transcription of SBE×4-luc under starvation (Fig. 7f). Homodeletion, which is the major form of genetic alterations of *ARL15* gene, and several nonsense mutations found in the cancer genomic database should render *ARL15* null and reduce the normal TGFβ signaling activity, as shown by our Arl15 knockdown experiment (Fig. 6b and Supplementary Fig. 6d). The TGFβ signaling pathway is tumor-suppressive for the proliferation of pre-malignant cancer cells(Hao et al., 2019; Lamouille et al., 2014; Massague, 2008; Seoane and Gomis, 2017). Therefore, our data demonstrate that cancer patients can have genetic mutations or alterations that compromise the Arl15-Smad4 interaction and suggest that such genetic changes might contribute to tumorigenesis by down-regulating the TGFβ signaling pathway.

## Discussion

Our study uncovered Arl15 as a unique regulator of the TGFβ family signaling pathway. According to our knowledge, it is the first small G protein reported to interact with Smads. It positively regulates both TGFβ and BMP pathways by directly interacting with the MH2 domain of common Smad, Smad4, and promoting the assembly of the Smad-complex. Interestingly, the Smad-complex serves as an effector and a GAP of Arl15 so that it dissociates Arl15 and subsequently enters the nucleus.

Our work provides a molecular mechanism on how closed or autoinhibited Smad4 becomes open or activated, an outstanding question in the field(Hata et al., 1997). We discovered that opening and, hence, activation of Smad4 is specifically aided by Arl15-GTP through direct binding to the MH2 domain of Smad4. A small G protein is regulated by its GEFs and GAPs, activities of which could be subject to further intracellular or extracellular stimuli. Hence, our study reveals Arl15 as a potential signaling integration node between a small G protein activation cascade and the TGFβ family signaling. The finding might provide us a new clue to understand the contextual and paradoxical nature of the TGFβ family signaling.

Based on our data, we propose a working model on the molecular role of Arl15 in the TGFβ family signaling (Fig. 7g). A currently unidentified GEF first activates Arl15-GDP to become GTP-loaded. Next, Smad4, mainly in a closed or inactive conformation, interacts with Arl15-GTP. Once bound to Arl15-GTP, Smad4-MH1 dissociates from its MH2 domain, rendering Smad4 in an open or active conformation. Though the Smad4-MH2 domain possesses a GAP activity toward Arl15, it might be too weak to inactivate Arl15-GTP *in vivo*. Therefore, a pool of Arl15-GTP-Smad4 complex might accumulate intracellularly. Upon stimulation by a TGFβ family cytokine, R-Smads (Smad2 as an example in Fig. 7g) are phosphorylated by TGFβ type I receptor kinase at their C-termini. With an activated MH2 domain, Arl15-bound Smad4 subsequently interacts with phospho-R-Smads. Since the R-Smad and Smad4 are known to assemble as a heterotrimer(Chacko et al., 2001; Chacko et al., 2004), it is tempting to speculate that one Arl15-GTP and one Smad-heterotrimer to form a heterotetramer. We further assume that associated Arl15 probably inhibits the nuclear translocation of the Smad-complex. By engaging R-Smads, the GAP activity of Smad4 is greatly enhanced and consequently triggers the GTP hydrolysis of Arl15. After the resulting Arl15-GDP dissociates from the Smad-complex, the latter translocates to the nucleus and executes eventual genomic actions.

Our model proposes that the Smad-complex is an effector and a terminator for active Arl15, which promotes the assembly of the Smad-complex in the first place. A similar case has been proposed for Sar1 and COPII coat complex(Antonny et al., 2001; Bi et al., 2002; Bi et al., 2007). Sar1 is an Arf-family small G protein that initiates the assembly of the COPII coat at the ER exit site. The COPII coat consists of repetitive units of two heterodimers, Sec23-Sec24, and Sec13-Sec31. In Sec23-Sec24 heterodimer, Sec23 functions as both an effector and a GAP for Sar1. The GAP activity of Sec23 is substantially boosted after Sec23-Sec24 recruits Sec13-Sec31 to complete the budding of a COPII-coated vesicle. The hydrolysis of Sar1-bound GTP eventually uncoats COPII-coated vesicles, which prepares them for the subsequent transport and fusion. We note that Arl15 and Sar1 have Ala (at 86) and His (at 77) in their G3-motifs, respectively, in contrast to Gln of other Arf-family small G proteins. Conserved Gln in the G3-motif functions to facilitate the GTP hydrolysis by correctly positioning the water molecule. Therefore, termination of active Sar1 and Arl15 *in vivo* probably requires their effector-cum-GAPs, which ensure the proper assembly of the COPII coat and the Smad-complex, respectively, before the complete dissociation of small G proteins. This property probably makes Sar1 and Arl15 a suitable initiator in promoting the assembly of large protein complexes. Confirmation of our speculation awaits future structural study of the Arl15-Smad-complex.

Our observation that Arl15 is essential for the invasion and migration of malignant cancer cells suggests that it might be a pro-metastatic factor. However, the finding of mutations compromising the Arl15-Smad4 interaction in the cancer genomic database suggests that Al15 might be a tumor suppressor. It is well known that the TGFβ signaling pathway possesses a context-dependent dual-role in cancer progression(Massague, 2008; Massague, 2012). In normal epithelia or pre-malignant cancers, the TGFβ signaling pathway exerts a cytostatic or tumor-suppressive effect to inhibit the proliferation of cells. On the other hand, in malignant or metastatic cancers, TGFβ signaling exerts a pro-metastatic effect to promote growth and invasion of cells.

An example is the effector of Arl15, Smad4(Deckers et al., 2006). Consistent with the tumor-suppressive function of Smad4, depletion of Smad4 promotes the growth of NMuMG mammary gland epithelial cells; in contrast, the depletion of Smad4 inhibits the metastasis of MDA-MB-231 cells, in agreement with the pro-metastatic role of Smad4. Furthermore, inactivating mutations are commonly found in the *SMAD4* gene in pre-malignant cancers such as pancreatic carcinoma, while *SMAD4* has been reported as an essential gene for metastasis of breast cancer(Levy and Hill, 2006; Massague, 2008; Massague, 2012). Therefore, we hypothesize that, like Smad4, Arl15 might play a similar dual-role in cancer progression.

Although our study mainly focused on the TGFβ signaling pathway, we observed that Arl15 promotes the assembly of the BMP Smad-complex (Supplementary Fig. 4), and it is essential for the BMP R-Smad-dependent transcription (Supplementary Fig. 6d). Therefore, our data indicate that Arl15 also participates in the BMP signaling pathway. Recent evidence suggests that TGFβ and BMP pathways play opposite roles in animal development and diseases(Ning et al., 2019). It has been proposed that competition between TGFβ and BMP pathways for Smad4 contributes to their antagonistic cellular effects(Candia et al., 1997; Sartori et al., 2013; Yuan et al., 2018). Hence, we speculate that, similar to Smad4, Arl15 might mediate antagonistic crosstalk between the two pathways.

Genetic studies have implicated the *ARL15* gene locus in rheumatoid arthritis and multiple metabolic diseases(Corre et al., 2018; Danila et al., 2013; Glessner et al., 2010; Li et al., 2014; Negi et al., 2013; Replication et al., 2014; Richards et al., 2009; Ried et al., 2016; Sun et al., 2015; Willer et al., 2013). Consistent with the broad role of the TGFβ signaling pathway in physiology and pathology, our findings raise the likelihood of *ARL15* as the causative gene and suggest the contribution of the TGFβ family signaling pathway to the pathology of these metabolic diseases. We hypothesize that the expression level of *ARL15* might vary in particular genetic background, therefore correspondingly changing the strength of the TGFβ family signaling. Indeed, we found that the overexpression and depletion of Arl15 substantially modulate the TGFβ-dependent transcription (Fig. 6a-d; Supplementary Fig. 6b,d,j,k). Therefore, our findings warrant a further investigation of the role of Arl15 in metabolic diseases.

## Materials and methods

### DNA plasmids

See Supplementary Table 1.

### Antibodies, TGFβ family cytokines, and chemicals

Mouse anti-Smad2/3 mAb (#610842; 1:1000 for Western blot or WB), mouse anti-GM130 mAb (#610823, 1:500 for immunofluorescence or IF) and mouse anti-Golgin245 mAb (#611280, 1:200 for IF) were purchased from BD Bioscience. Rabbit anti-phospho-Smad2 (S465/467)/Smad3 (S423/425) mAb (#8828; 1:1000 for WB, 1: 200 for IF) and rabbit anti-Flag mAb (#14793; 1:1000 for WB) were purchased from Cell Signaling Technology. Rabbit anti-Nup133 mAb (#ab155990; 1:1000 for WB) was purchased from Abcam. The following antibodies were from Santa Cruz: mouse anti-Smad4 mAb (B-8, #sc-7966,1:1000 for WB), rabbit anti-Smad1/5/8 polyclonal antibody (pAb) (#sc-6031-R, 1:1000 for WB), rabbit anti-glyceraldehyde 3-phosphate dehydrogenase (GAPDH) pAb (#sc-25778, 1:1000 for WB), mouse anti-GFP mAb (#sc-9996, 1:1000 for WB, 1: 200 for IF), mouse anti-His mAb (#sc-8036, 1:1000 for WB), mouse anti-Myc mAb (#sc-40, 1:1000 for WB, 1:200 for IF) and mouse anti-HA mAb (#sc-7396, 1:1000 for WB). HRP (horseradish peroxidase)-conjugated goat anti-mouse (#176516, 1:10,000 for WB) and anti-rabbit IgG antibodies (#176515, 1:10,000 for WB) were from Bio-Rad. Alexa Fluor conjugated goat anti-mouse (1:500 for IF), anti-rabbit IgG antibodies (1:500 for IF) and recombinant human BMP2 (#PHC7145) were from Thermo Fisher Scientific. Recombinant human TGFβ1 (#100-21C-10) was purchased from PeproTech. 20 μg ml^-1^ stock solution of TGFβ1 was made in 4 mM HCl containing 1 mg ml^-1^ bovine serum albumin. SB431542 (#1614) and guanosine 5′-[β,γ-imido]triphosphate (GMPPNP, #G0635) were purchased from Tocris and Sigma-Aldrich, respectively.

### Cell culture and transfection

HeLa, HEK293T, MCF7, and MDA-MB-231 cells were from American Type Culture Collection. 293FT cells were from Thermo Fisher Scientific. MCF7, MDA-MB-231, HeLa, HEK293T and 293FT cells were maintained in high glucose Dulbecco’s Modified Eagle’s Medium (DMEM) supplemented with 10% fetal bovine serum (FBS) (Thermo Fisher Scientific) at 37 °C in a 5% CO_2_ incubator. FBS was heat-inactivated at 55 °C for 30 min. Cells were transfected using polyethylenimine (Polysciences) or Lipofectamine 2000 transfection reagent (Thermo Fisher Scientific).

During live-cell imaging, transfected HeLa cells grown on a glass-bottom Petri-dish (MatTek Corporation) were imaged in the CO_2_-independent medium (Thermo Fisher Scientific) supplemented with 4 mM glutamine and 10% FBS at 37 °C.

### Lentivirus-mediated knockdown and expression

293FT cells grown in a 6-well plate were transfected with shRNA constructs in pLKO.1 vector or pLVX expression constructs together with pLP1, pLP2, and pLP/VSVG using Lipofectamine 2000 (Thermo Fisher Scientific). 18 h after transfection, cells were incubated with fresh medium (DMEM supplemented with 10% FBS) at 37 °C for another 24-48 h. The supernatant of the tissue culture medium was collected, passed through a 0.45 μm filter (Sartorius), and used immediately. For lentivirus-mediated knockdown or expression, cells were incubated for 24-48 h with the virus supernatant supplemented with 8 µg ml^-1^ polybrene (Sigma-Aldrich #H9268) before being subjected to further experimental procedures.

### Preparation of recombinant proteins

DNA plasmids encoding recombinant proteins were used to transform BL21 *E coli* bacterial cells. After induction by isopropyl β-D-1-thiogalactopyranoside, a bacterial cell pellet was collected. For purification of GST-fusion proteins (those cloned in pGEB or pGEX-KG vectors), cells were lysed using the freeze-thaw method in a buffer containing 50 mM Tris pH 8.0, 150 mM NaCl, 2 mM dithiothreitol (DTT), 1 mg ml^-1^ lysozyme and 1 mM phenylmethylsulfonyl fluoride (PMSF). The cell lysate was centrifuged at 20,000 g for 30 min, and the resulting supernatant was incubated with Glutathione Sepharose 4B beads (GE Healthcare) overnight at 4 °C. The beads were washed three times with a buffer containing 50 mM Tris pH 8.0, 0.1% Triton-X 100, 150 mM NaCl and 2 mM DTT. The bound proteins were eluted using reduced glutathione, and the resulting eluent was subjected to extensive dialysis against phosphate-buffered saline (PBS).

For purification of His-tagged proteins (those cloned in pET30a and pET30ax vectors), bacterial cells were lysed with a buffer containing 20 mM Tris pH 8.0, 150 mM NaCl, 10 mM imidazole, 2 mM DTT, 1 mg ml^-1^ lysozyme, and 1 mM PMSF and the resulting supernatant was incubated with nickel-nitrilotriacetic acid agarose (QIAGEN) overnight at 4 °C. The beads were washed with a buffer containing 20 mM Tris pH 8.0, 150 mM NaCl, 20 mM imidazole, and 2 mM DTT and eluted with an elution buffer containing 20 mM Tris pH 8.0, 150 mM NaCl, 250 mM imidazole, and 2 mM DTT. The eluent was subjected to extensive dialysis against PBS. All purified proteins were quantified by Coomassie staining in SDS-PAGE.

### Production of Arl15 antibody

DNA plasmid, His-Arl15-WT in pET30ax, was used to transform BL21 *E coli* cells. After induction by isopropyl β-D-1-thiogalactopyranoside, the bacterial cell pellet was lysed by sonication in PBS containing 8 M urea. After high-speed centrifugation, the supernatant was incubated with nickel-nitrilotriacetic acid agarose beads at room temperature for 2 h. The beads were subsequently washed in PBS containing 8 M urea and 20 mM imidazole. Next, bead-bound His-Arl15-WT was eluted in PBS containing 8 M urea and 250 mM imidazole. After concentrating and changing the buffer to PBS containing 4 M urea, His-Arl15-WT was sent to Genemed Synthesis Inc for rabbit immunization and antiserum collection. To purify the polyclonal antibody against Arl15, GST-Arl15 immobilized on glutathione Sepharose beads was incubated with 50 mM dimethyl pimelimidate (Thermo Fisher Scientific) in 200 mM sodium borate pH 9.0 to crosslink GST-Arl15 covalently onto glutathione Sepharose beads. The crosslinked beads were subsequently washed with 200 mM ethanolamine pH 8.0, incubated with the antiserum at room temperature for 1 h and washed by PBS. Finally, the bound antibody was eluted by 100 mM glycine pH 2.8 and dialyzed against PBS.

### Co-IP and GST pull-down

HEK293T cells transfected by indicated DNA constructs were lysed with the lysis buffer (40 mM HEPES pH 7.4, 150 mM NaCl, 1% Trition X-100, 2.5 mM MgCl_2_, 1 × cOmplete™ Protease Inhibitor Cocktail (Roche), 1 mM PMSF, and 1 mM DTT). For co-IP, after centrifugation, the supernatant was incubated with indicated antibodies overnight at 4 °C. The antibody-antigen complex was captured by incubating with lysis buffer prewashed proteinA/G beads (Thermo Fisher Scientific) for 2 h. In some co-IPs, GFP-Trap beads (ChromoTek) were used to directly IP GFP-tagged fusion proteins. For GST pull-down, cleared cell lysate was incubated with bead-immobilized GST-fusion protein for 4 - 14 h at 4 °C. The beads were subsequently washed with the lysis buffer. Next, the bound protein was eluted by boiling in SDS-sample buffer and resolved in the SDS-PAGE. SDS-PAGE separated proteins were transferred to polyvinyl difluoride membrane (Bio-Rad), which was sequentially incubated with the primary and HRP-conjugated secondary antibody. At last, the chemiluminescence signal was detected by a cooled charge-coupled device using LAS-4000 (GE Healthcare Life Sciences). Alternatively, the chemiluminescence signal was detected by CL-XPosure^TM^ film (Thermo Fisher Scientific) and digitally scanned.

### GTP-agarose pull-down

Transiently transfected HEK293T cells were suspended in a binding buffer (20 mM HEPES, pH7.4, 150 mM NaCl, 1 × cOmplete™ Protease Inhibitor Cocktail). After three cycles of freeze-thaw and extrusion through a 25 ½ gauge needle, the cell lysate was cleared by centrifugation at 16,000 g for 30 min and subsequently treated with 5 mM EDTA (final concentration) for 1 h at 4 °C. Next, the lysate was incubated with GTP-agarose beads (bioWORLD) for 1 h in the presence of 10 mM MgCl_2_ (final concentration) at 4 °C. After washing with the binding buffer four times, beads were boiled in the SDS-sample buffer and analyzed by Western blot.

### IF

Cells grown on Φ12 mm glass coverslips were fixed with 4% paraformaldehyde in PBS at room temperature for 20 min and washed with 100 mM ammonium chloride and PBS. Next, cells were sequentially incubated with primary and fluorescence-conjugated secondary antibodies, which were diluted in the fluorescence dilution buffer (PBS supplemented with 5% FBS and 2% bovine serum albumin) containing 0.1% saponin (Sigma-Aldrich). After extensive washing with PBS, coverslips were mounted in the Mowiol mounting medium, containing 12% Mowiol 4-88 (EMD Millipore), 30% glycerol, and 100 mM Tris pH 8.5.

### Wide-field fluorescence microscopy

Unless specified, all fluorescence images were acquired under an inverted wide-field fluorescence microscope (Olympus IX83) equipped with a Plan Apo oil objective lens (63× or 100× oil, NA 1.40), a motorized stage, motorized filter cubes, a scientific complementary metal oxide semiconductor camera (Neo; Andor Technology), and a 200 watt metal-halide excitation light source (LumenPro 200; Prior Scientific). Dichroic mirrors and filters in filter cubes were optimized for Alexa Fluor 488/GFP, 594/mCherry and 647. The microscope system was controlled by MetaMorph software (Molecular Devices), and only the center quadrant of the camera sensor was used for imaging. During the live-cell imaging, HeLa cells grown on a glass-bottom Petri-dish (as described above in Cell culture and transfection) were imaged in a 37 °C chamber.

### Laser scanning confocal microscopy

HeLa cells co-expressing mCherry and GFP-tagged proteins were grown on a glass-bottom Petri-dish as described above (Cell culture and transfection). Live-cell imaging was conducted in a 37°C chamber under Zeiss LSM710 laser scanning confocal microscope system (Carl Zeiss) equipped with a Plan-apochromat objective (100x oil, NA 1.40). Two laser lines with wavelengths of 488 nm and 561 nm were used to excite GFP and mCherry, and their emission filter bandwidths were 495-550 nm and 595-620 nm, respectively. The microscope system was controlled by ZEN software (Carl Zeiss).

### Dual-luciferase assay

HeLa cells cultured in 24-well plates were transfected with indicated firefly luciferase reporter construct (SBE×4-luc or BRE-luc) together with pRL-SV40 renilla luciferase control vector (Promega) using Lipofectamine 2000 (Thermo Fisher Scientific). Constant amount of total transfected DNA was balanced by supplying pBluescript SK vector DNA to the transfection mixture. 24 h after transfection, cells were serum-starved for 4 h and treated with 5 ng ml^-1^ TGFβ1 for 20 h. Cells were subsequently lysed, and firefly and renilla luciferase activities were measured using the Dual-Luciferase Reporter Assay System (Promega) according to standard protocol.

### RT-qPCR

Total RNA was extracted from MCF7 or MDA-MB-231 cells using Trizol^TM^ reagent (Thermo Fisher Scientific) according to the manufacturer’s protocol. Reverse transcription primed by random nonamer primers was conducted using nanoScript 2 Reverse Transcription kit (Primerdesign). The qPCR was performed using SYBR green based PrecisionFAST kit (Primerdesign) in Bio-Rad CFX96 Touch™ real-time PCR detection system. Melt curves and agarose gel electrophoresis were performed to confirm the specificity of PCR primers. The qPCR result of each gene was first divided by that of β-tubulin and further normalized to that of control (empty vector or GL2-shRNA treatment). Primers for qPCR are as follows: c-Myc (5’-AAAGGCCCCCAAGGTAGTTA-3’; 5’-GCACAAGAGTTCCGTAGCTG-3’), ID1 (5’-CAAATTCAAGGTGGAATCGAA-3’; 5’-GGTGGCTGGGAAGTGAACTA-3’), p21^cip1^ (5’-GAGGCCGGGATGAGTTGGGAGGAG-3’; 5’-CAGCCGGCGTTTGGAGTGGTAGAA-3’), p27^kip1^ (5’-GCTCCACAGAACCGGCATTT-3’; 5’-AAGCGACCTGCAACCGACGATTCTT-3’), E-cadherin (5’-TCTTCCCCGCCCTGCCAATC-3’; 5’-GCCTCTCTCGAGTCCCCTAG-3’), N-cadherin (5’-GGTGGAGGAGAAGAAGACCAG-3’; 5’-GGCATCAGGCTCCACAGT-3’), vimentin (5’-CTAGGAGCCCTCAATCGG-3’; 5’-CACGGACCTGGTGGACAT-3’), β-tubulin (5’-TTGGCCAGATCTTTAGACCAGACAAC-3’; 5’-CCGTACCACATCCAGGACAGAATC-3’) and Arl15 (5’-CCCCGATAACGTCGTGTC-3’; 5’-AGCGGCTCCAGTATTTCC-3’).

There were some modifications in the experiment described in Fig. 6d. Lentivirus was harvested by transfecting pMD2.G, psPAX2 together with Arl15-shRNA1, Arl15-shRNA2 or GL2-shRNA in pLKO.1. After lentivirus infection, pooled MCF7 cells were transiently selected with puromycin. The resulting cells were treated with 5 ng ml^-1^ TGFβ1 for 72 h and total RNA was extracted by using TRIzol^TM^ reagent and Qiagen RNeasy Mini Kit (Qiagen) according to the manufacturer’s protocol. Reverse transcription primed by random hexamer primer was conducted using RevertAid H Minus First Strand cDNA Synthesis Kit (Thermo Fisher Scientific). SensiFAST™ SYBR® Hi-ROX Kit (Bioline) in QuantStudio™ 5 (Thermo Fisher Scientific). The qPCR result of each gene was first divided by that of ribosomal protein L13A mRNA (Primers: 5’-GCC TTC ACA GCG TAC GA-3’; 5’-CGA AGA TGG CGG AGG TG-3’) and further normalized to that of GL2-shRNA control.

### Nuclear fractionation

5 x 10^6^ HeLa cells collected by scraping culture flasks were washed three times with ice-cold PBS. After centrifugation at 200 g for 5 min, pelleted cells were re-suspended in 500 μl buffer A (20 mM Tris pH 7.4, 10 mM NaCl, 3 mM MgCl_2_, 0.5 % NP40, 1 mM DTT and 1 mM PMSF) and incubated on ice for 15 min. Cells were subsequently vortexed for 10 sec, and the resulting cell lysate was centrifuged for 10 min at 1,000 g at 4 °C. The supernatant, which the non-nuclear or cytoplasmic fraction, was transferred into a new tube, while the pellet, which is the nuclear fraction, was washed three times using buffer A without NP40. Both fractions were subjected to the SDS-PAGE and Western blot analysis.

### Image analysis

All image analysis was conducted in ImageJ (http://imagej.nih.gov/ij/).

### GAP assay

A similar protocol has been previously described(Pan et al., 2006). Briefly, purified His-Arl15-WT and His-Arl15-AL proteins were incubated with a 20-fold molar excess of GTP in 20 mM HEPES pH7.5, 150 mM NaCl, 5 mM ethylenediaminetetraacetic acid, and 1mM DTT at room temperature for 1 h. Next, the proteins were subjected to a 7 KDa molecular weight cut-off Zeba Spin Desalting Column (Thermo Fisher Scientific), which was pre-equilibrated with 20 mM HEPES pH7.5 and 150 mM NaCl. The GAP assay was conducted in a 96-well glass-bottom microplate (Corning), and the released inorganic phosphate was measured using EnzChek Phosphate Assay Kit (Thermo Fisher Scientific). The reaction system contained 20 mM HEPES pH 7.5, 150 mM NaCl, 0.15 mM 2-amino-6-mercapto-7-methylpurine ribonucleoside, 0.75 U ml^-1^ purine nucleoside phosphorylase, 10 mM MgCl_2_, 40 μM GTP-loaded His-Arl15-WT or His-Arl15-AL, and 0.4 µM following GAP candidate proteins, single or in combinations as indicated in text: His-Smad4, His-Smad2-SE, GST-Smad4, GST-Smad4-MH1, GST-Smad4-linker-MH2, GST-Smad4-MH2, and GST (negative control). The kinetics of the GTP hydrolysis was continuously monitored by the absorbance at 360 nm in Cytation 5 (BioTek) at 22 °C. For each time series, absorbance values were subtracted by the corresponding initial value measured at 0 min.

### ER-to-Golgi and Golgi export trafficking assays

These assays were performed as previously described(Mahajan et al., 2019). HeLa cells subjected to lentivirus-transduced shRNA knockdown were further transfected to express a RUSH reporter: Ii-Strep_ManII-SBP-GFP or Ii-Strep_TNFα-SBP-GFP(Boncompain et al., 2012) for the ER-to-Golgi or Golgi export to the PM transport assay. 50 ng ml^-1^ streptavidin was added to cell culture and was removed 20 h after transfection of RUSH reporters. For the ER-to-Golgi trafficking assay, 40 µM biotin and 10 µg ml^-1^ cycloheximide were added to the cell medium during the chase. For the Golgi export trafficking assay, cells were first incubated at 20 °C for 3 h in the presence of 40 µM biotin and 10 µg ml^-1^ cycloheximide to accumulate TNFα-SBP-GFP at the Golgi. The system was subsequently warmed up to 37 °C during the chase. In both assays, cells were processed for immunofluorescence at various chase times and imaged by the wide-field microscope. The Golgi fraction of RUSH reporter is calculated using *I*_Golgi_/*I*_cell_, in which *I*_Golgi_ and *I*_cell_ are integrated GFP intensity of the Golgi and the cell, respectively. All cells positively expressing RUSH reporter were analyzed in each image.

### Cancer mutation data of Arl15

Cancer mutation data of Arl15 were manually compiled from cBioPortal (https://www.cbioportal.org/) and COSMIC (https://cancer.sanger.ac.uk/cosmic/gene/analysis?ln=ARL15).

### Wound healing migration assay

MDA-MB-231 cells were first subjected to lentivirus-transduced knockdown using GL2-shRNA, Arl15-shRNA1, or 2. Next, cells were cultured to confluence in 6-well plates. Gaps were scratched across a well using a pipette tip, and the closure of gaps was kinetically monitored by an inverted phase contrast microscope. The widths of gaps were quantified using ImageJ (https://imagej.nih.gov/ij/). The percentage of the relative migration was calculated as (1-*d*/*d_0_*)*100%, in which *d_0_* is the initial width of the gap and *d* is the width of the gap at a specific time.

### Invasion assay

MDA-MB-231 cells were first subjected to lentivirus-transduced knockdown using GL2, Arl15-shRNA1, or 2. After cells were serum-starved for 24 h in DMEM, the same amount of suspended cells were added into upper chambers of Corning Costar Transwell cell culture inserts (pore size 8 µm; Sigma-Aldrich, #CLS3464) with the coating of 30 μg Matrigel^TM^ Basement Membrane Matrix (BD Biosciences). 700 µl DMEM supplemented with 10 % FBS was added into the lower chamber. After 24 h incubation at 37°C in a CO_2_ incubator, the upper chamber was washed three times with PBS, fixed with ice-cold methanol, and stained with crystal violet solution (0.5 % crystal violet in 20 % methanol). Finally, cells on the upper surface of the insert were removed with a cotton swab, and those on the lower surface (translocated cells) were imaged. In each experiment, five randomly selected fields were imaged, and the number of cells within each image was counted and averaged. The relative invasion was calculated as the number of cells per image normalized by that of control (GL2-shRNA treated).

## Author contributions

LL conceived the study. LL and LV supervised the study. LL and MS designed experiments. MS conducted the majority of experiments. HCT performed the experiments for Fig. 1a,b, and Supplementary Fig. 1b,c. HCT and XS contributed to Fig. 2 experiments. MD did the experiments for Supplementary Fig. 6e-h. YZ identified Smad4 as a potential Arl15 effector via the yeast-two hybrid screening. YZ and BKH confirmed the interaction between Arl15 and Smad4. MS and LL analyzed data and prepared figures. LL wrote the manuscript.

## Acknowledgments

We would like to thank Z. Ding (Temasek Polytechnic, Singapore) for the help in the yeast-two hybrid screening and R. Derynck (University of California San Francisco, USA), P. ten Dijke (Leids Universitair Medisch Centrum, Netherlands), W. Hong (Institute of Molecular and Cell Biology, Singapore), T. Kirchhausen (Harvard Medical School, USA), M. Lowe (University of Manchester, UK), F. Perez (Institut Curie, France), D. Trono (EPFL, Switzerland), D. Root (Broad Institute, USA), M. Roussel (St. Jude Children’s Research Hospital, USA) for sharing DNA plasmids. This work was supported by the following grants to L.L.: MOE AcRF Tier1 RG35/17, Tier2 MOE2015-T2-2-073, and MOE2018-T2-2-026.

## List of Supplementary Materials

Supplementary Table 1

Supplementary Fig. 1-7

## Supplementary Information

### Supplementary figure legends

**Supplementary Figure 1.**
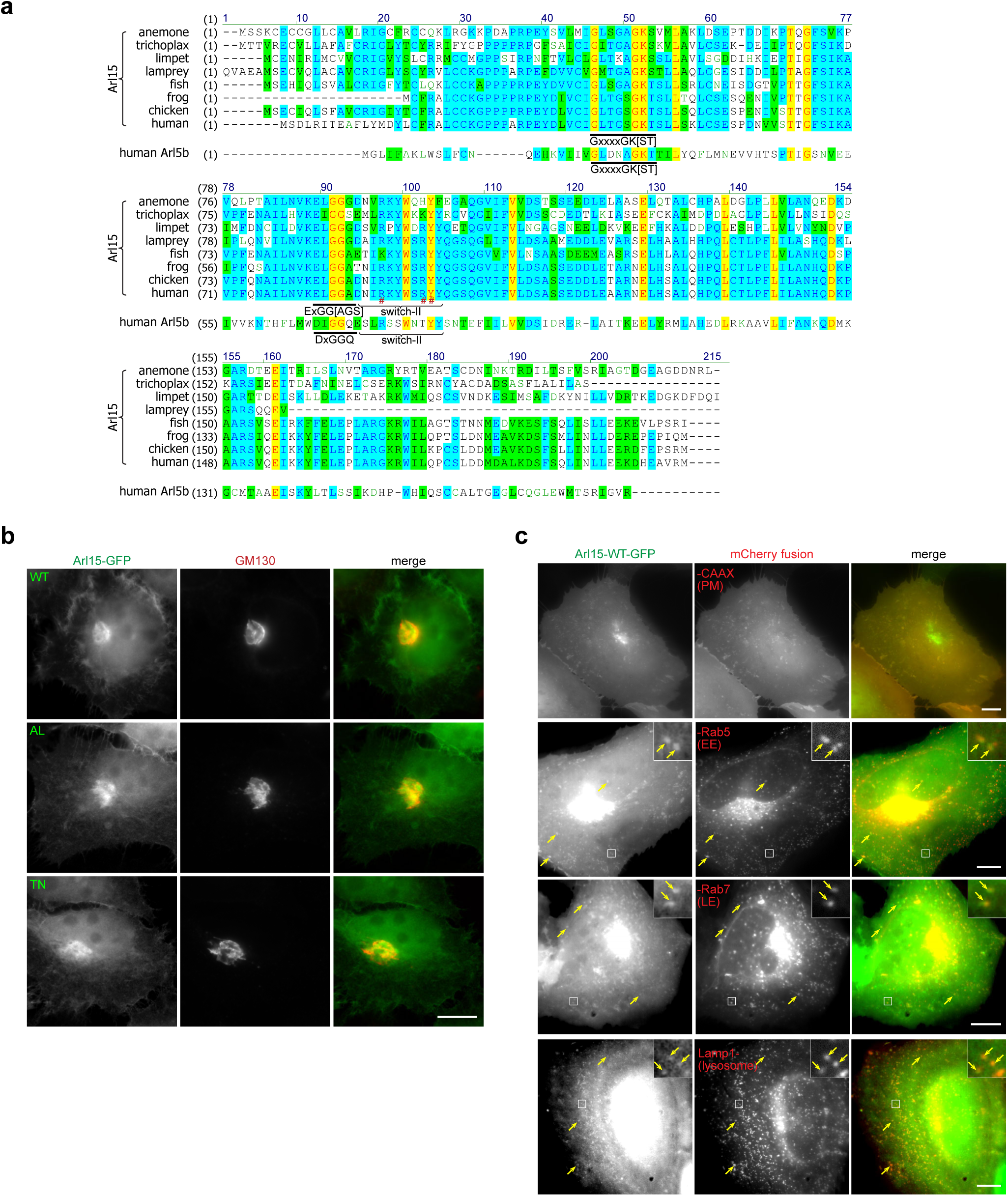
Multiple sequence alignment of Arl15 and Arl5b and sub-cellular localization of Arl15. **a** Multiple sequence alignment of metazoan Arl15 orthologues and human Arl5b. The conserved G1 (GxxxxGK[ST]) and G3 (ExGG[AGS]) motifs (DxGGQ for Arl5b) are underlined, and the switch-II region is marked. In human Arl15 sequence, R90, R95, and Y96 are indicated by #. The following UniProt sequences of Arl15 were used: anemone, A7SYP3; trichoplax, B3RRR0; limpet, V4BVB7; lamprey, S4R642; fish, A5PMK4; chicken, F1NMW9; frog, F6WKV0; human, Q9NXU5. The Uniprot ID of human Arl5b is Q96KC2. **b** Arl15-WT, AL, and TN mutants localize to the Golgi. HeLa cells transiently expressing Arl15-(WT, AL, or TN)-GFP were immunostained for endogenous GM130. **c** C-terminally GFP-tagged Arl15 can be detected at the PM (mCherry-CAAX), EE (mCherry-Rab5), LE (mCherry-Rab7), and lysosome (Lamp1-mCherry). HeLa cells transiently co-expressing Arl15-WT-GFP and indicated mCherry-tagged organelle marker were imaged live. Boxed regions are zoomed in in the upper right corners to show colocalization. Arrows indicate colocalization. All images were acquired by a wide-field microscope. Scale bar, 10 µm.

**Supplementary Figure 2.**
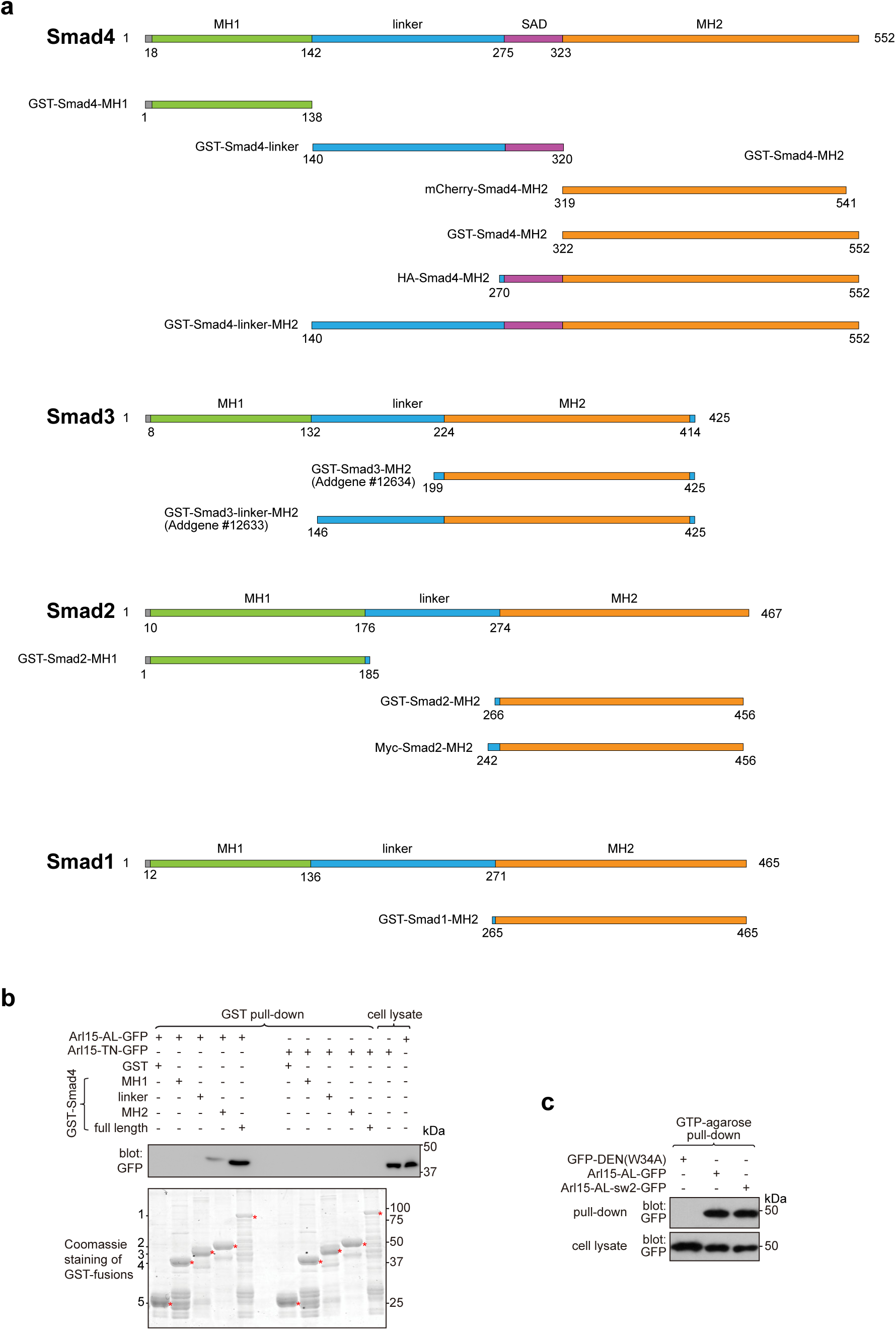
A schematic diagram showing the domain organization of Smad constructs, a pull-down assay investigating the interaction between Arl15 and Smad4 domains, and a GTP-binding assay demonstrating the proper folding of Arl15 switch-II mutant. **a** A schematic diagram showing the domain organization of Smad1, 2, 3, and 4 constructs. Numbers indicate corresponding amino acid positions in the full-length protein sequence. SAD, Smad4 activation domain. **b** The linker region of Smad4 does not interact with the GTP-mutant form of Arl15. Bead-immobilized GST-fusions of Smad4 fragments were incubated with HEK293T cell lysate expressing Arl15-AL-GFP or Arl15-TN-GFP, and pull-downs were immunoblotted against GFP. Loading of GST-fusions was shown below by Coomassie staining. 1, GST-Smad4; 2, GST-Smad4-MH2; 3, GST-Smad4-linker; 4, GST-Smad4-MH1; 5, GST. * indicates specific band. **c** Arl15 switch-II mutant, Arl15-AL-sw2-GFP, can bind to GTP. HEK293T cells lysates expressing indicated GFP-tagged proteins were incubated with the GTP-agarose, and pull-downs were immunoblotted against GFP. GFP-DEN(W34A), which is not a G protein, serves as a negative control. In **b** and **c**, molecular weights (in kDa) are labeled in all immunoblots.

**Supplementary Figure 3.**
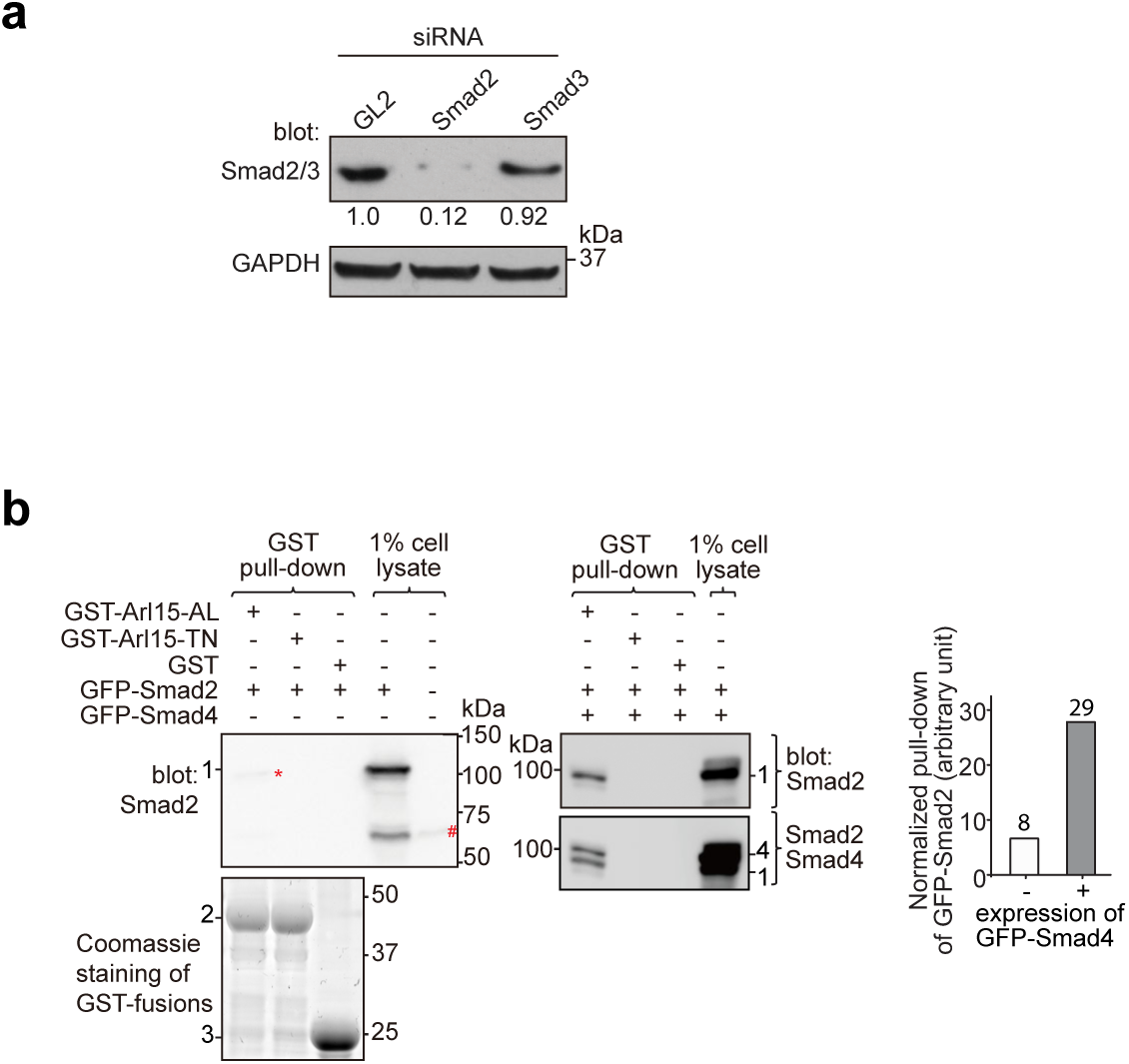
Arl15-GTP indirectly interacts with Smad2 via Smad4. HEK293T cells under the normal culture condition were used. **a** The anti-Smad2/3 mAb primarily detects endogenous Smad2 in HEK293T cells. Cells were subjected to GL2, Smad2, or Smad3 siRNA knockdown, and the resulting cell lysates were blotted for indicated proteins. Ratios of the intensity of Smad2/3 band to that of GAPDH band are displayed below. **b** The GTP-mutant form of Arl15 pulls down more exogenously expressed Smad2 when Smad4 was co-expressed. Bead-immobilized GST-fusion proteins were incubated with cell lysates expressing indicated proteins, and pull-downs and 1% cell lysate inputs were immunoblotted against Smad2 (anti-Smad2/3 mAb) and Smad4. In the middle panel, the same blot was sequentially blotted for Smad2 (upper blot) followed by Smad4 (lower blot). 1, GFP-Smad2; 2, GST-Arl15 (AL or TN); 3, GST; 4, GFP-Smad4; #, endogenous Smad2; *, the weak band of GFP-Smad2 that was pulled down without co-expression of Smad4. The normalized pull-down of GFP-Smad2, calculated by the ratio of the intensity of the pull-down band to that of 1% cell lysate input band, is plotted in the right panel. Loading of fusion proteins is shown by Coomassie staining. Molecular weights (in kDa) are labeled in immunoblots and gels.

**Supplementary Figure 4.**
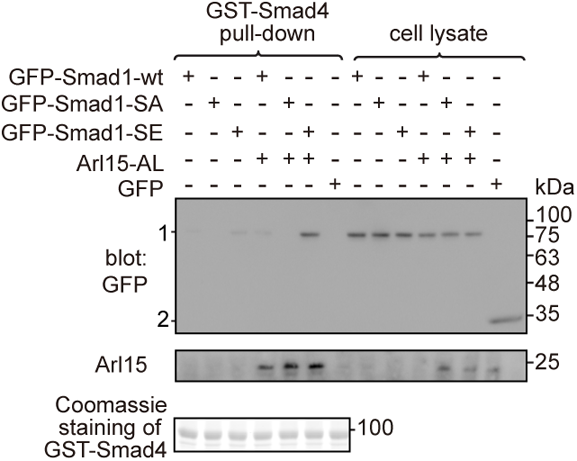
Arl15-GTP promotes the interaction between Smad4 and phospho-Smad1. HEK293T cells were cultured under normal condition. Bead-immobilized GST-Smad4 was incubated with cell lysates expressing indicated proteins, and pull-downs and the cell lysates were immunoblotted for GFP or Arl15. 1 and 2 indicate GFP-Smad1 (WT, SA, or SE) and GFP bands, respectively. Loading of fusion proteins is shown by Coomassie staining. Molecular weights (in kDa) are labeled in the immunoblot.

**Supplementary Figure 5.**
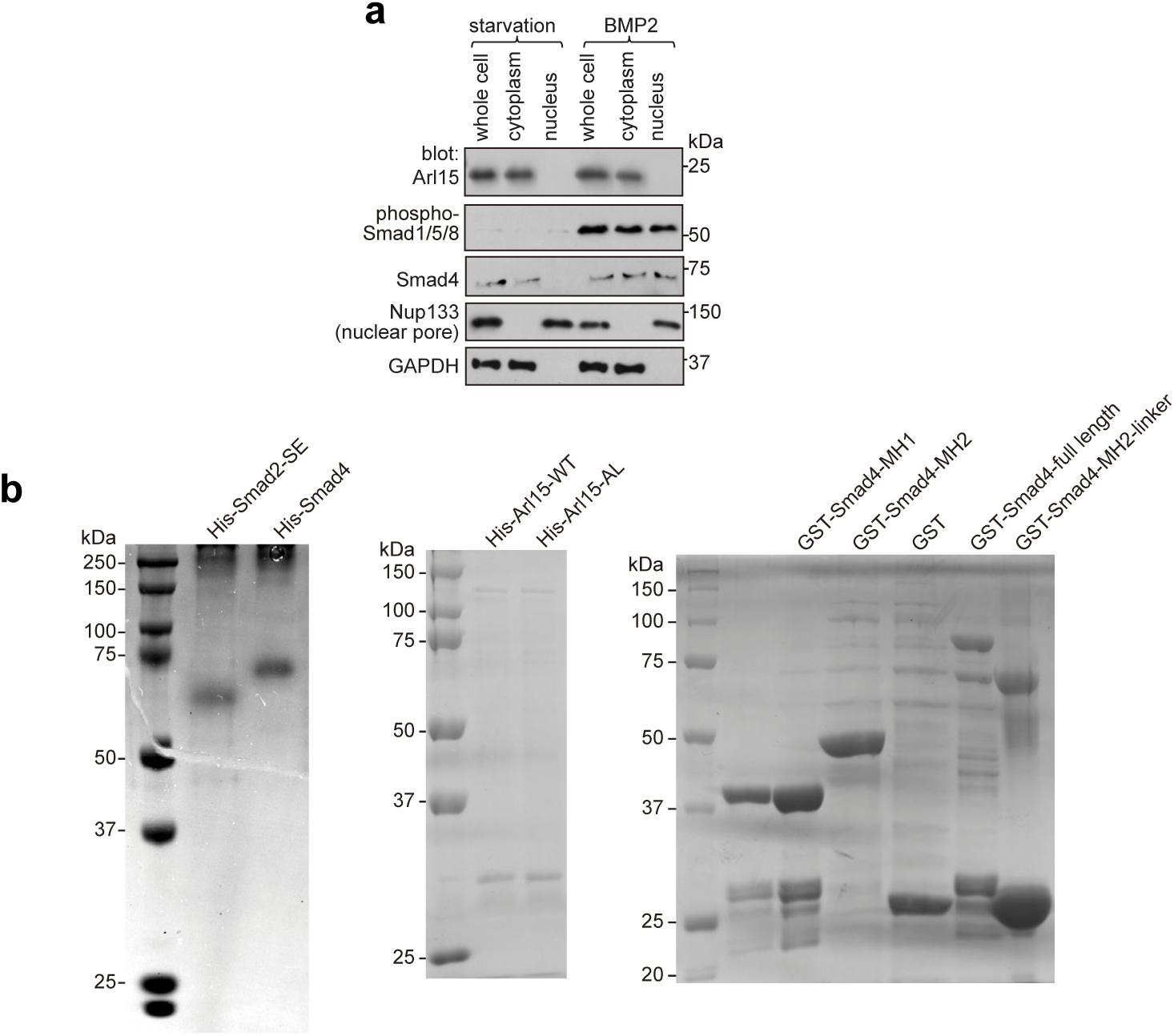
Nucleus translocation of phospho-Smad1/5/8 and Coomassie gels of purified proteins used in GAP assays. **a** Under BMP2 treatment, phospho-Smad1/5/8, but not Arl15, translocates to the nucleus. As described in Figure 5b, serum-starved HeLa cells were either further serum-starved or treated with 100 ng ml^-1^ BMP2 for 4 h. Total cell lysate and nuclear and cytosol fractions were subjected to immunoblotting against indicated proteins. **b** Coomassie gels of purified proteins used in the GAP assay described in Figure 5d,e. Molecular weights (in kDa) are labeled in all immunoblots and gels.

**Supplementary Figure 6.**
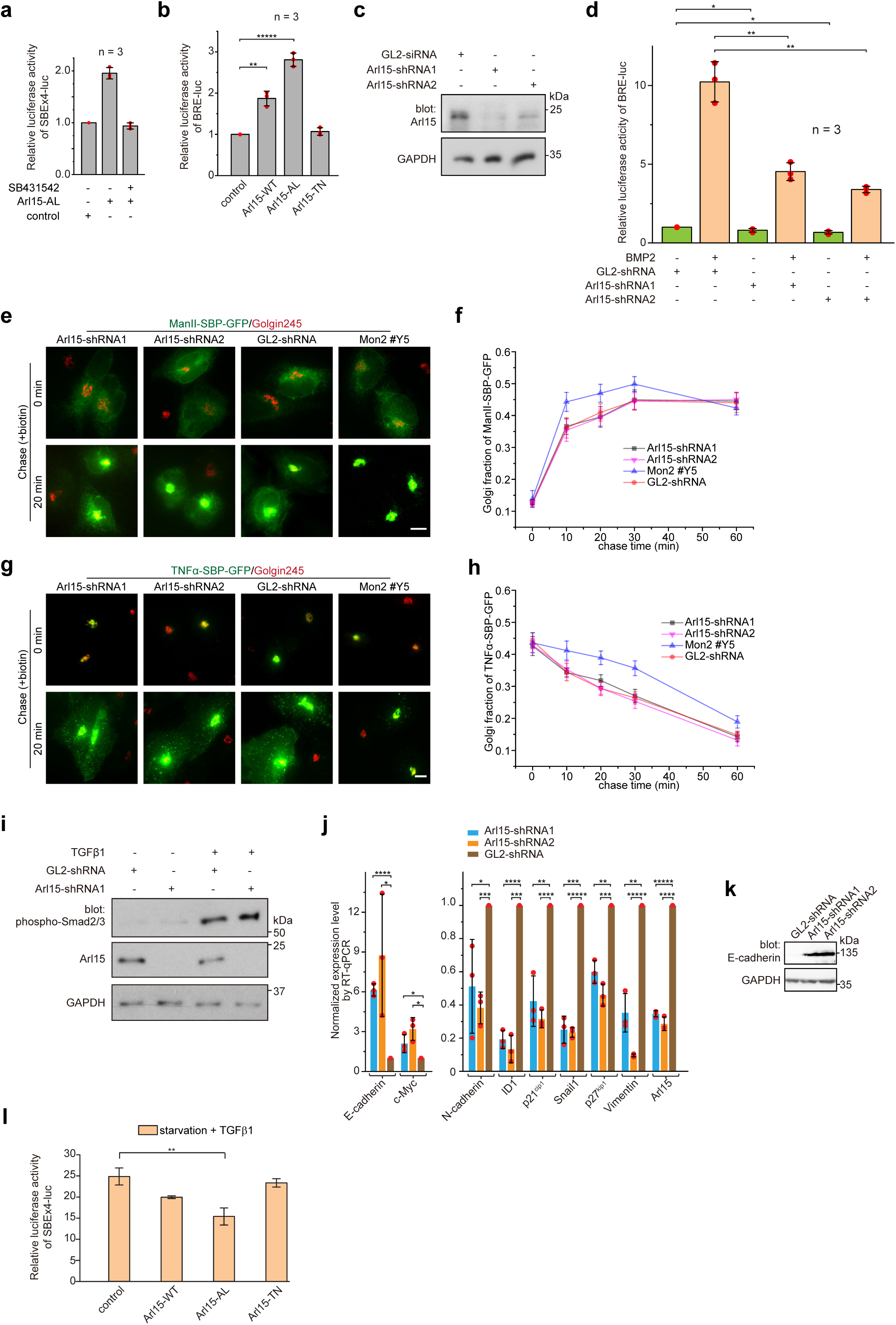
Arl15 promotes the TGFβ family signaling pathway. **a** The Arl15-AL- induced transcription of SBE×4-luc requires the kinase activity of the TGFβ type I receptor. HeLa cells co-expressing SBE×4-driven firefly luciferase and SV40-driven renilla luciferase together with Arl15-AL or pBluescript SK vector DNA (control) were serum-starved for 4 h followed by 20 h treatment with starvation medium with or without 2 μM SB431542. The relative luciferase activities were subsequently acquired and normalized. **b** Arl15 positively regulates the BMP signaling pathway since overexpression of Arl15-WT or AL, but not TN, promotes the transcription of BRE-luc reporter. HeLa cells co-expressing BRE-driven firefly luciferase and SV40-driven renilla luciferase together with indicated Arl15 mutant, or pBluescript SK vector DNA (control) were serum-starved for 24 h. The relative luciferase activities were subsequently acquired and normalized. **c** Endogenous Arl15 can be depleted by lentivirus-transduced shRNA knockdown. HeLa cells were lysed after lentivirus-transduced knockdown using indicated shRNA, and the resulting cell lysates were immunoblotted for Arl15 and GAPDH. **d** Arl15 is essential for efficient BMP signaling since its depletion reduces BMP2-stimulated transcription of BRE-luc reporter. After lentivirus-transduced knockdown of Arl15, HeLa cells expressing the dual-luciferase as described in **b** were serum-starved for 4 h followed by 20 h treatment with starvation medium supplemented with or without 100 ng ml^-1^ BMP2. The relative luciferase activities were subsequently acquired and normalized. **e, f** The ER-to-Golgi transport of ManII does not require Arl15. In **e**, after lentivirus-transduced knockdown of Arl15, HeLa cells transiently expressing RUSH reporter ManII-SBP-GFP were subjected to biotin treatment to chase ManII-SBP-GFP along the ER-to-Golgi pathway (see Materials and methods). Golgi fractions were plotted in **f**. **g**,**h** The Golgi export of TNFα does not require Arl15. In **g**, after lentivirus-transduced knockdown of Arl15, HeLa cells transiently expressing RUSH reporter TNFα-SBP-GFP was incubated at 20 °C in the presence of biotin to accumulate TNFα-SBP-GFP at the Golgi. Cells were subsequently incubated at 37 °C to chase the reporter to exit the Golgi. The Golgi fractions were plotted in **h**. Mon2 knockdown (Mon2 #Y5) serves as a control as it can accelerate the ER-to-Golgi transport and delay the Golgi export of secretory cargos. Scale bar, 10 µm. Error bar, mean ± SEM of *n* ≥ 30 cells. **i** Arl15 is not required for TGFβ1-stimulated phosphorylation of Smad2/3. After lentivirus-transduced knockdown of Arl15, HeLa cells were serum-starved for 4h followed by 20 h treatment with starvation medium supplemented with or without 5 ng ml-1 TGFβ1. Cell lysates were subsequently blotted for indicated proteins. **j** Depletion of Arl15 counteracts TGFβ1-induced transcriptions – it downregulates the transcription of N-cadherin, ID1, Snail1, vimentin, p27^kip1^, and p21^cip1^, and upregulates the transcription of E-cadherin and c-Myc. After lentivirus-transduced knockdown of Arl15, MDA-MB-231 cells were subjected to 16 h starvation followed by 5 ng ml^-1^ TGFβ1 treatment for 4 h. Transcripts of indicated genes were quantified and normalized as in Figure 6d. **k** Depletion of Arl15 upregulates the expression of E-cadherin in MDA-MB-231 cells. Knockdown was conducted as in **d**. Cell lysates were immunoblotted for E-cadherin and GAPDH. In **c**, **i**, and **k,** molecular weights (in kDa) are labeled in immunoblots. **l** Arl15-AL inhibits TGFβ1-induced transcription of SBE×4-luc. The experiment was conducted as described in Figure 6a except that all cells were treated with 5 ng ml^-1^ TGFβ1. In **a**, **b**, **d**, **j**, and **l,** error bar, mean ± SD of *n* = 3 experiments. *p* values were from the *t*-test (unpaired and two-tailed). NS, not significant (*p* > 0.05); *, *p* ≤ 0.05; **, *p* ≤ 0.005; ***, *p* ≤ 0.0005; ****, *p* ≤ 0.00005; *****, *p* ≤ 0.000005. Red dot, individual data point.

**Supplementary Fig. 7.**
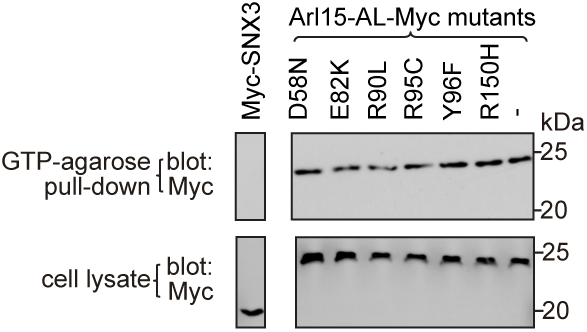
Arl15 missense mutations identified from cancer patients do not affect the GTP-binding of Arl15-AL. The figure corresponds to Figure 7e,f. HEK293T cell lysates transiently expressing indicated Myc-tagged proteins were incubated with the GTP-agarose, and pull-downs were immunoblotted against Myc. Myc-SNX3, which is not a G protein, serves as a negative control. Molecular weights (in kDa) are labeled. Bots in the same row are cropped from the same gel blot.

**Supplementary Table 1.**
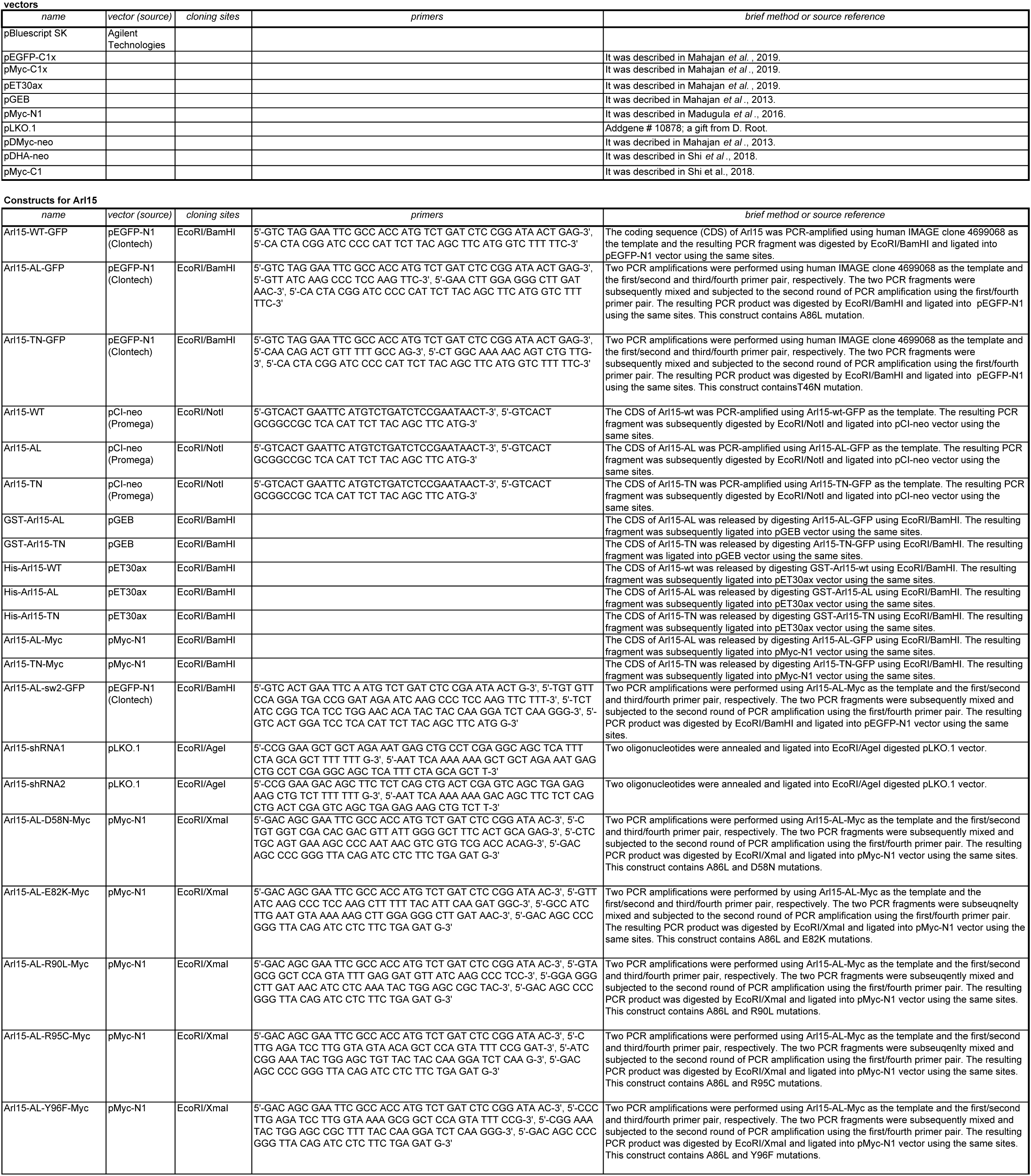
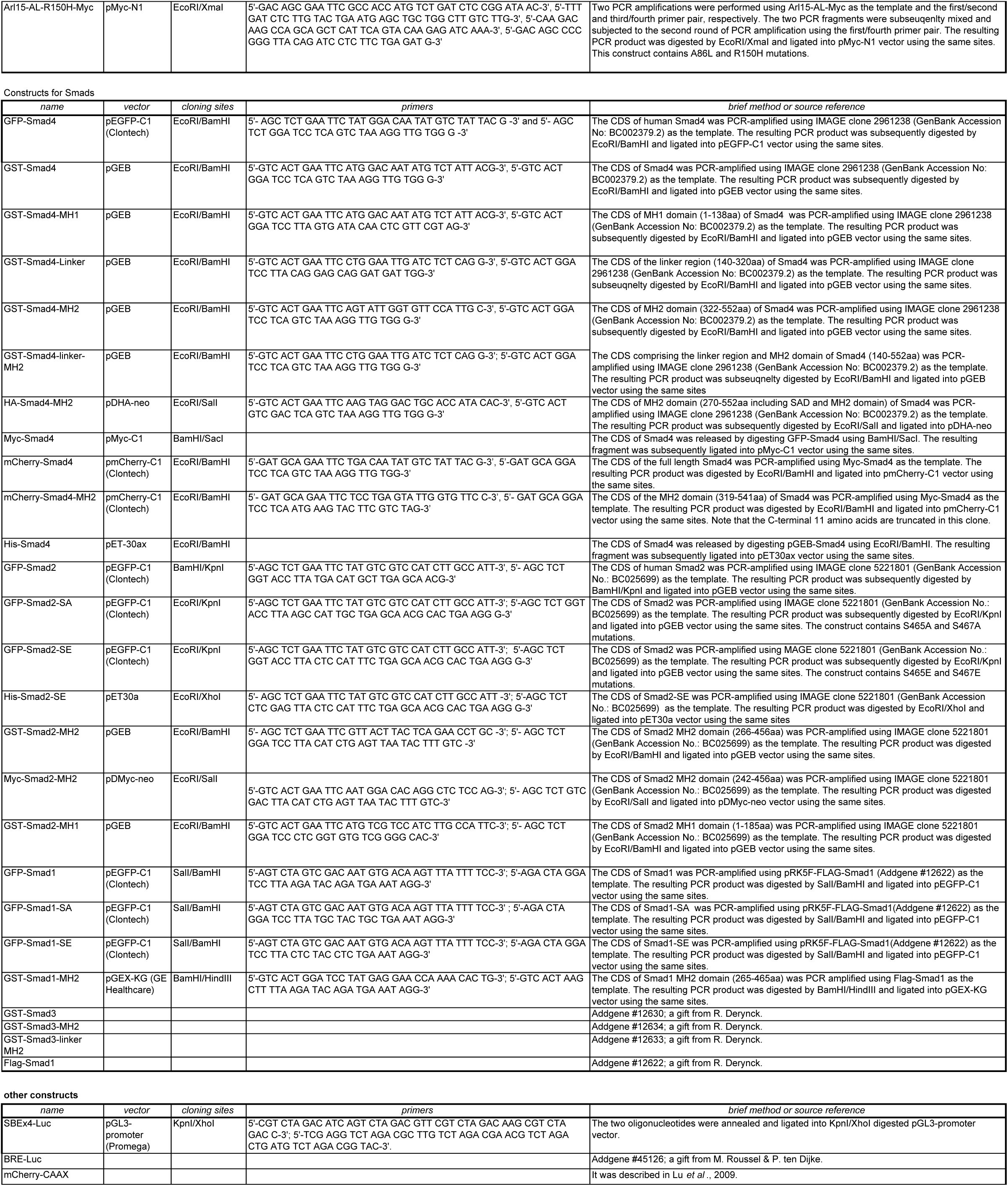
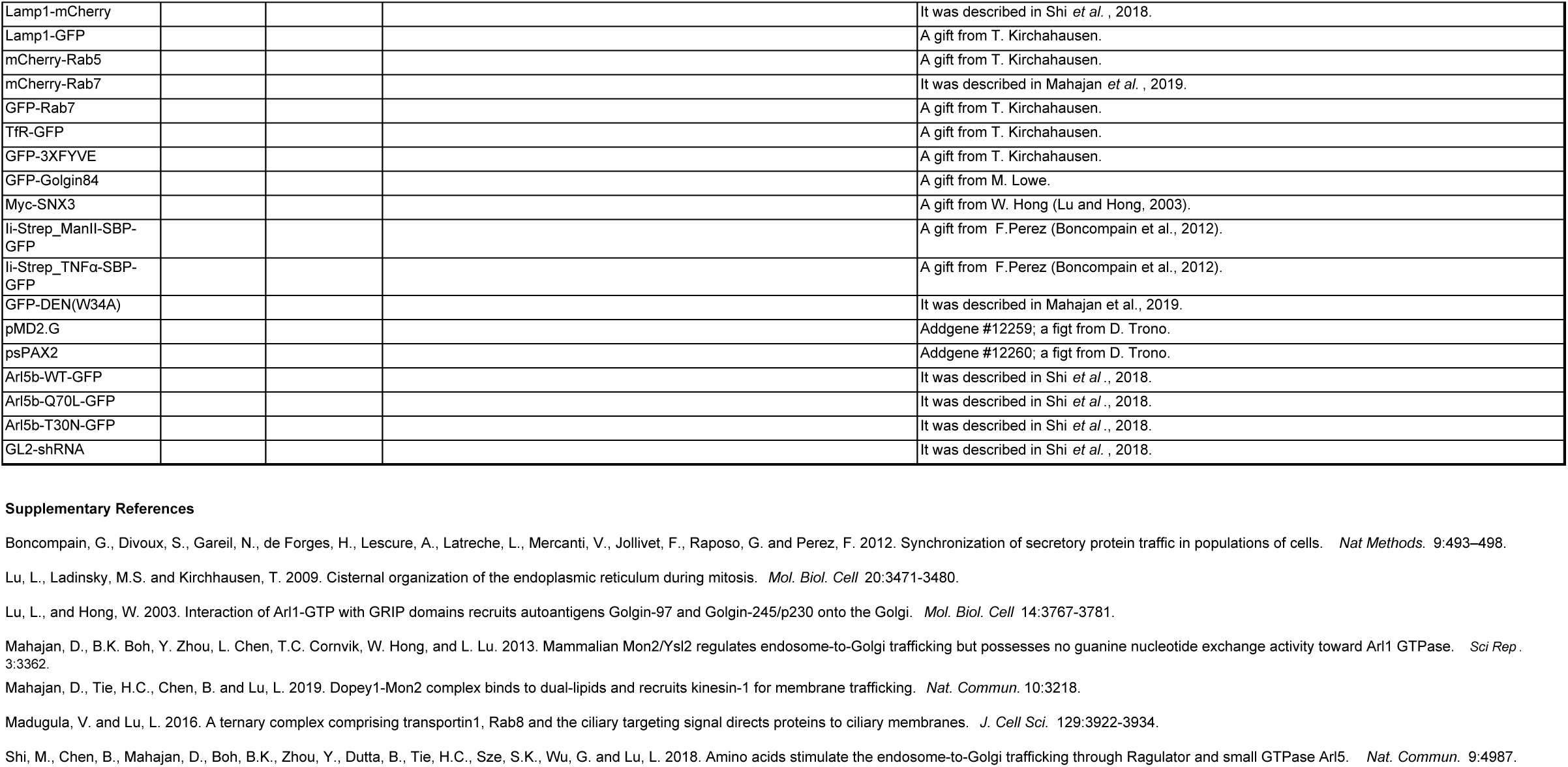
List of DNA plasmids used in this study.

## Notes

### Competing Interest Statement

The authors have declared no competing interest.

